# Comparative cross-species transcriptomic analysis identifies new candidates of Pooideae nitrate response

**DOI:** 10.64898/2026.03.18.712634

**Authors:** Mathilde Grégoire, Stéphanie Pateyron, Véronique Brunaud, Jean Philippe Tamby, Lamia Benghelima, Marie-Laure Martin, Thomas Girin

## Abstract

Nitrogen fertilizers are essential for crop productivity but cause environmental harm, necessitating the development of cultivars that thrive under limited nitrogen. This study investigates the transcriptomic response to nitrate in *Arabidopsis thaliana* (a model dicot), *Brachypodium distachyon* (a model Pooideae), and *Hordeum vulgare* (barley, a domesticated Pooideae) to identify conserved and species-specific molecular mechanisms. Using RNA-seq after 1.5 and 3 hours of nitrate treatment, we found that core nitrate-responsive biological processes – such as nitrate transport, assimilation, carbon metabolism, and hormone signaling – are largely conserved across species. However, comparative analysis at gene level based on orthology revealed specificities between the species. For instance, rRNA processing was uniquely stimulated in Arabidopsis, while cysteine biosynthesis from serine and gibberellin biosynthesis were specifically regulated in Brachypodium and barley. Orthologs of key nitrate-responsive genes (e.g., *NRT*, *NLP*, *TCP20*) exhibited variable regulation, reflecting potential adaptations linked to domestication or nutrient acquisition strategies. These findings highlight the importance of integrating model and crop species to uncover targets for improving nitrogen use efficiency in cereals. This study provides a general pipeline combining Gene Ontology enrichment and orthology analysis to compare transcriptomic responses across species. This workflow is highly useful for any comparative transcriptomic study in which differences in annotation quality and gene divergence between genomes may bias interpretation.

## Introduction

Pooideae, a sub-family of Poaceae, includes major temperate C3 cereals like barley (*Hordeum vulgare*), wheat (*Triticum sp.*), rye (*Secale cereale*) and oat (*Avena sativa*), which are vital for human and livestock nutrition. Brachypodium (*Brachypodium distachyon*) serves as a model for this sub-family (Girin *et al*., 2014). As the global population increases, the demand for food rises, making optimal crop production a critical challenge. Nitrogen (N) is a critical nutrient for plant growth, mainly absorbed as nitrate (NO₃⁻) and ammonium (NH₄⁺). While N fertilizers enhance agricultural productivity, their excessive use is costly and leads to environmental issues, including eutrophication and biodiversity loss. A main challenge of today’s agriculture is to maintain or enhance crop production while diminishing polluting fertilizer amounts. However, current cultivars have been bred under high nutrient availability, limiting their efficiency in low-N environments. Understanding the molecular mechanisms of nitrate response in both bred and wild species is essential for developing crops that thrive under limited N availability.

Nitrate is the primary N source for many plants, including Arabidopsis (*Arabidopsis thaliana*) and Pooideae crops. It is transported from the soil by NRT transporters and assimilated through enzymatic pathways involving nitrate reductase (NR), nitrite reductase (NiR) and the GS/GOGAT (Glutamine Synthetase/Glutamate Synthase) cycle (Xu *et al*., 2012). Nitrate also acts as a signaling molecule, regulating gene expression, growth, and development (Krouk *et al*., 2010).

In *Arabidopsis*, the Primary Nitrate Response (PNR) has been extensively studied, revealing key molecular actors. NRT1.1, a dual-affinity nitrate transporter, and NLP7, a transcription factor, function as nitrate sensors (Ho *et al*., 2009; Liu *et al*., 2022). Upon nitrate perception, PNR is rapidly induced by calcium signals perceived by CPKs which in turn phosphorylate NLP7 (Liu *et al*., 2017). Phosphorylated NLP7 is retained in the nucleus and physically interacts with NLP6, TCP20 and NRG2 to govern expression of genes necessary for nitrate metabolism (Marchive *et al*., 2013; Xu *et al*., 2016; Guan *et al*., 2017). Additional transcription factors, such as TGA1 and TGA4 (Alvarez *et al*., 2014), HRS1 (Medici *et al*., 2015; Maeda *et al*., 2018) and LBD37/38/39 (Rubin *et al*., 2009; Brooks *et al*., 2019) also play critical roles in nitrate signaling. Nitrate-responsive processes have been identified, including nitrate transport and assimilation, carbon metabolism, hormone signaling (particularly those involved in modulating root system architecture), and amino acid biosynthesis (Scheible et al., 2004; Canales et al., 2014; Ruffel et al., 2025).

Previous cross-species analyses between Arabidopsis and rice (Obertello *et al*., 2015) or between maize and sorghum (Du *et al*., 2020) identified conserved nitrate-responsive modules and targets that might be specific to monocots. A hormone-related module was identified, which could be linked to differences in root architecture between monocots and dicots (Smith and de Smet, 2012), thus to a divergence of its regulation by nitrate.

This study aimed to compare nitrate response in Arabidopsis (*Arabidopsis thaliana*; a wild dicot), Brachypodium (*Brachypodium distachyon*; a wild Pooideae) and barley (*Hordeum vulgare*; a domesticated Pooideae). We performed an RNAseq analysis on roots in response to 1.5 and 3 hours of 1mM nitrate treatment, preceded by 4 days of N starvation. We then compared their responses using gene ontology (GO) enrichment to estimate conservation or divergence of the responses in terms of biological processes. To deepen the comparison at the gene level, we performed a global orthology analysis. We thus identified common regulated processes such as global nitrate transport and assimilation system as well as hormone-related processes. However, at gene level within most of studied processes, we identified differential regulation between species. Finally, combining GO enrichments and OGps allowed us to identify processes specifically regulated in one or two species only, such as the rRNA pathway – specifically upregulated in Arabidopsis – or the cysteine biosynthesis from serine pathway – specifically upregulated in Brachypodium and barley.

## Materials and methods

### Plant growth conditions

Three independent replicates were produced, with 1 week between plant batches. N-deprived basal medium (0.35mM K_2_SO_4_, 0.25mM CaCl_2_, 0.25mM KH_2_PO_4_, 0.25mM MgSO_4_, 10 mg.L^-1^ Fer-EDTA, 243 µM Mo_7_O_2_4(NH_4_)_6_, 0.4 µM H_3_BO_3_, 118 µM SO_4_Mn, 10 µM SO_4_Cu, 34.8 µM SO_4_Zn) was adapted from Castle and Randall (1987). Arabidopsis (Col-0) seeds were sown on gelled basal medium in 1.5 mL cut tubes (0.65% Plant Agar; pH 5.8) supplemented with 0.05mM KNO_3_ and 0.025mM Ca(NO_3_)_2_. Tubes were placed on hydroponics systems containing liquid basal medium supplemented with 0,05mM KNO_3_ and 0,025mM Ca(NO_3_)_2_ in a growth chamber (16h/8h day/night, 21°C/18°C ; 180 µE/m^2^/s). Medium was renewed after 7, 11 and 14 days. Brachypodium (Bd21-3) and barley (Golden Promise) seeds were stratified in water for 3 days at 4°C, then sown on petri dishes with water at room temperature in the dark. After 7 days, plantlets were placed on hydroponics systems containing liquid basal medium supplemented with 0.05mM KNO_3_ and 0.025mM Ca(NO_3_)_2_ in a growth chamber (18h/6h day/night, 22°C/19°C ; 250 µE/m^2^/s). Medium was renewed after 4 and 7 days. Eighteen days after sowing, Arabidopsis, Brachypodium and barley plantlets were transferred to N-deprived basal medium supplemented with 5.5mM KCl. Medium was renewed after 3 days. Twenty-two days after sowing, plantlets were transferred to basal medium supplemented with either 1mM NO_3_^-^ (treated; 0.5mM KNO_3_, 0.25mM Ca(NO_3_)_2_) or 5.5mM KCl (mock). Roots of 4 plants per sample were harvested after 1.5h or 3h of treatment and frozen in liquid nitrogen.

### RNA extraction

Fifty mg of frozen root tissue were frozen-ground, resuspended in 500 µL NucleoZol and 200 µL RNase-free H_2_O and incubated for 30min at room temperature. Tubes were vortexed and incubated for 30min at room temperature, then centrifuged for 15min à 4°C (12000 RCF). Supernatants were collected and 2.5 µL of 4-bromoanisole were added before vortexing. Samples were centrifuged 10min at 4°C (12000 RCF) and 500µL of supernatant were collected in new tubes. 500µL of Propane-2-ol were added before 40min incubation at room temperature. After 25min centrifugation (12000 RCF) at room temperature, pellets were washed twice with 75% ethanol. Pellets were air-dried and resuspended with 50µL RNase-free water. Samples were then purified using Clean and concentrator (Zymo Research) following the manufacturer’s instructions. RNA concentrations were quantified using a Nanodrop.

### RT-qPCR on sentinels’ genes

cDNA were synthesized using 0.7 µg of RNA, oligo(dT)_18_ and RevertAid H Minus Reverse Transcriptase kit (Thermo Scientific), following the manufacturer’ instructions, and diluted 1:20. For qPCR, 2.5 µL cDNA were used with 2.35 SYBR mix (SsoAdvanced™ Universal SYBR® Green Supermix, Bio-Rad) and 0.03 µM of forward and reverse primers in a final volume of 10 µL. Relative quantification and statistics were performed following Rieu and Powers (2009). Normalization was done by dividing Ct of genes of interest by Ct of a synthetic gene, the geometric mean of 2 (barley) or 3 (Arabidopsis and Brachypodium) housekeeping gene Cts. Primers for qPCR experiments are listed in Supp Table 1.

### RNA-seq technology

Quality of total RNA was controlled on an Agilent 2100 Bioanalyzer, according to the manufacturer’s recommendations. Library construction was performed by the IPS2-POPS platform (Gif-sur-Yvette, France). Briefly, mRNAs were polyA selected, fragmented to 260 bases and libraries were built using the TruSeq stranded mRNA kit (Illumina®, California, U.S.A.), with an Applied BioSystem 2720 Thermal Cycler and barcoded adaptors. Barcoded libraries were sequenced on an Illumina NexSeq500 sequencer (IPS2 POPS platform). RNA-seq samples were sequenced in single-ends, stranded mode with a sizing of 260bp and a read length of 75 bases, lane repartition and barcoding giving approximately 20-40 million reads per sample.

### RNA-seq bioinformatics treatments and analyses

RNA-seq preprocessing included trimming library adapters and performing quality controls. Raw data (fastq) were trimmed with Trimmomatic (Bolger *et al*., 2014) for Phred Quality Score >20 and read length >30 bases. Ribosomal sequences were removed with sortMeRNA (Kopylova *et al*., 2012). The genomic mapper STAR v2.7 (Dobin *et al*., 2013) was used to align reads against the genomes with options --outSAMprimaryFlag AllBestScore and --outFilterMultimapScoreRange 0 to keep the best results. The three genomes and annotation files used were *Hordeum vulgare* v1 (IBSC), *Brachypodium distachyon Bd21.3* v1 (Phytozome) and *Arabidopsis thaliana* v10 (TAIR). Gene abundance was calculated with STAR, counting only reads mapping unambiguously to one gene and removing multi-hits. According to these rules, 94% of reads were associated with a unique gene for Arabidopsis and Brachypodium, and 88% for barley.

Differential analyses were performed independently for each species, following Rigaill *et al*. (2018). Briefly, genes with less than 1 read after count-per-million (CPM) normalization in at least half of the samples were discarded. Library size was normalized using the trimmed mean of M-value (TMM) method and count distribution was modeled with a negative binomial generalized linear model. The logarithm of the mean expression was defined as a function of a time effect, a treatment effect and their interaction. Dispersion was estimated by the edgeR method (version 3.28.0; McCarthy *et al*., 2012) in the statistical software ‘R’ (version 3.6.1 R; Development Core Team; 2005). Quality control identified a failed sample (Bd_Mock_3h_2), so biological repeat #2 of the 3h mock treatment was removed for differential analysis for all three species. Expression differences of interest were tested using a likelihood ratio test and p-values were adjusted by the Benjamini-Hochberg procedure to control False Discovery Rate (FDR). A gene was declared differentially expressed if its adjusted p-value < 0.05.

### Data Deposition

All steps of the experiment, from growth conditions to bioinformatics analyses, were managed in CATdb database (Gagnot *et al*., 2008; http://tools.ips2.u-psud.fr/CATdb/) with ProjectID NGS2019_07_NiCe. This project was submitted from CATdb into the international repository GEO (Gene Expression Omnibus; Edgar *et al*., 2002; http://www.ncbi.nlm.nih.gov/geo), access ID GSE317477.

### Orthology analysis

Amino-acid sequences of proteomes were collected as FASTA files, based on annotations TAIR v10 for *Arabidopsis thaliana*, Bd21-3 v1 for *Brachypodium distachyon* and High Confidence v2 for *Hordeum vulgare*. Only representative protein forms were used (103 797 protein sequences), excluding alternative forms and multiple variants. Orthologous genes were investigated using OrthoFinder (v2.3.7; Emms and Kelly, 2015). The first step of OrthoFinder (all-versus-all sequence comparisons) was performed with Diamond (v0.9.24; Buchfink *et al*., 2015). All subsequent steps were managed by OrthoFinder to produce orthogroups (OGps) and identify orthologs.

### Gene ontology analysis

Biological process Gene Ontology analyses were performed using the topGO R package (Alexa *et al*., 2006; Alexa and Rahnenführer, 2016) with the elim algorithm and Fisher’s exact test, with R version 3.6.3. GO databases from EnsemblPlants were used for *Arabidopsis thaliana*, *Brachypodium distachyon* and *Hordeum vulgare*.

The Normalized Enrichment Score (NES) was calculated as the relative enrichment of a GO in a species *versus* the enrichment in all three species: NES_speciesX = [ES_speciesX/(ES_species1+ES_species2+ES_species3)]x100, with ES_speciesX = (number_DEGs_in_GOterm_speciesX/number_detected_genes_in_GOterm_speciesX)).

## Results

### Effect of 1mM Nitrate treatment on Arabidopsis, Brachypodium and barley transcriptomes

The goal of this study was to identify common and specific processes, genes and gene groups responding to nitrate in the three species by transcriptomics. To identify the maximum responding genes, harvesting was done at two timepoints after nitrate treatment, limiting biases due to possible differences in response kinetics between species. Experimental conditions were adjusted for Brachypodium to maximize nitrate response (data not shown) and validated on the other species. Three independent nitrate-treatment experiments were performed, each containing one biological replicate. Plantlets were cultivated 18 days on basal medium with 0.1mM NO_3_^-^, before transfer to N-deprived medium for 4 days, then to medium containing 1mM NO_3_^-^ or no N source for 1.5h or 3h (Supp Figure 1). Roots were collected for RNA extraction, RT-qPCR and RNAseq analyses. Expression of sentinel genes *NRT2.1*/*NRT2A*, *NIA1* and *NIA2* was significantly induced after both 1.5h and 3h nitrate treatment (except *HvNIA2*, induced only after 1.5h; Supp Figure 2), validating experimental conditions.

RNA-seq read mapping on genomes led to unambiguous detection of 19 498 genes in Arabidopsis, 20 845 in Brachypodium and 19 076 in barley (66%, 57% and 48% of annotated genes, respectively; Supp Table 2). Comparing nitrate-treated to mock samples, 3 888, 3 658 and 4 440 Differentially Expressed Genes (DEGs) were identified respectively across both timepoints (Supp Figure 1; Supp Table 2). For each timepoint-species combination, upregulated and downregulated gene numbers were similar (Supp Figure 3A-C). Many DEGs responded at both timepoints (34–53% of total induced or repressed DEGs). DEG numbers were higher at 3h than 1.5h (increase of 8–33%, considering stimulated and repressed genes separately; Supp Figure 3A-C), suggesting a response amplification between timepoints. In agreement, genes responding at both timepoints globally showed a stronger response at 3 h (Supp Figure 3D-F). Five Arabidopsis, 4 Brachypodium and 16 barley genes showed opposite regulation at 1.5 h *vs*. 3 h (Supp Figure 3; Supp Table 2).

To identify biological processes associated with DEGs per species, Gene Ontology (GO) enrichment analysis was performed (Figure 1). TopGO R package was used, as it accounts for GO hierarchy thus enables the identification of most relevant GO terms. Analysis used the full DEG sets (*ie* combination of 1.5h and 3h sets) for all DEGs, upregulated only or downregulated only (Figure 2A; Supp Table 3A). Separate 1.5h and 3h analyses gave similar results (Supp Figures 4A & 5A; Supp Tables 3B & 3C), suggesting a unique response phase in terms of biological processes over the time period.

**Figure 1:**
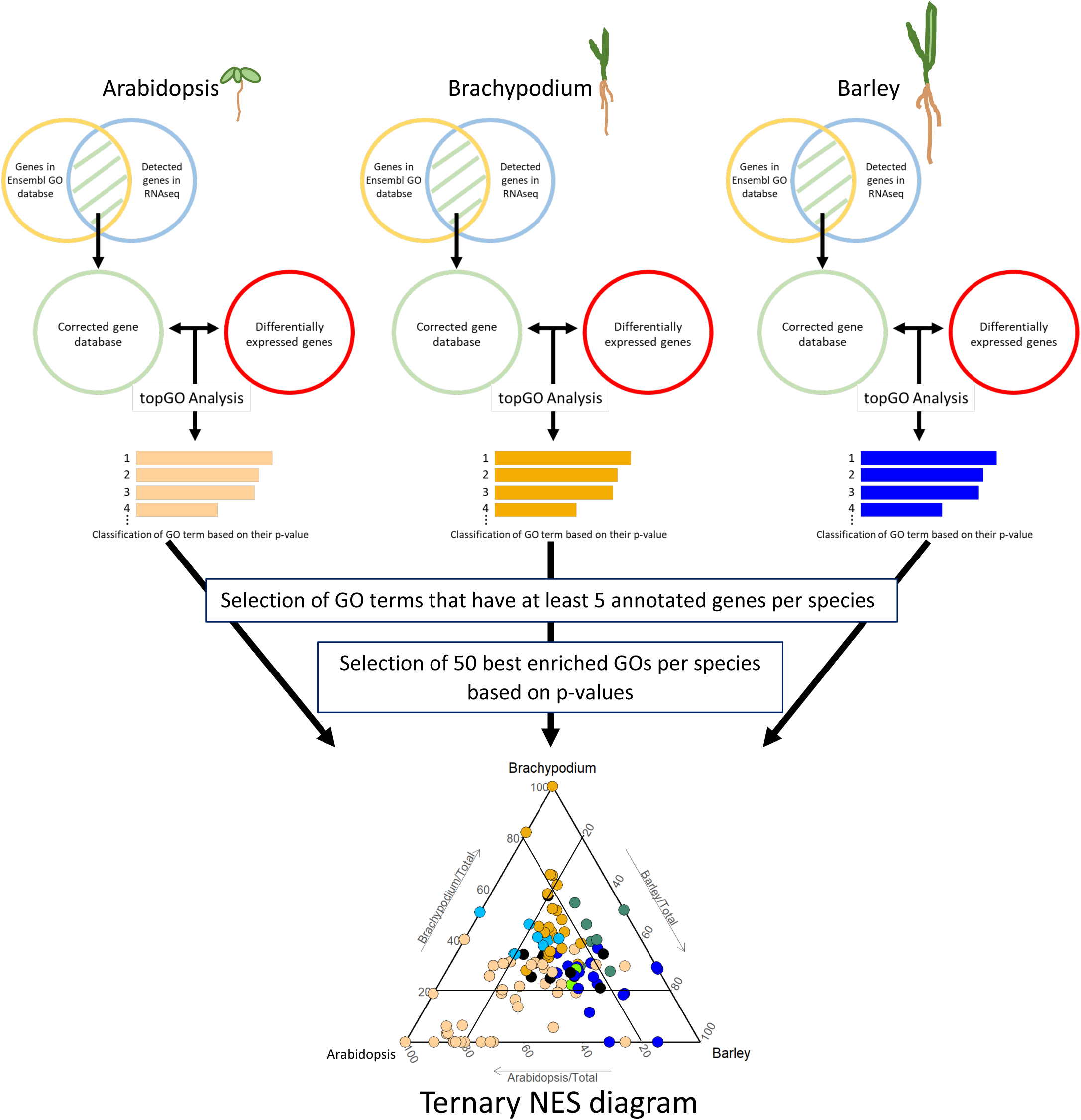
Schematic diagram of the Gene ontology enrichment analysis. Ensembl databases were filtered to detected genes only. Differentially expressed gene lists were analyzed using the topGO R package to rank GO terms by p-value (elim Fisher’s exact test). To minimize annotation biases between species, only GO terms containing ≥5 detected genes per species were retained. The 50 most significantly enriched GO terms per species were selected for comparative analysis. Normalized Enrichment Scores (NES; see Materials and Methods) were calculated per GO-species combination and plotted as ternary diagrams. Each dot represents a GO term; triangle corners indicate maximum NES for the species (GO terms with no regulated genes in other species); internal bars represent thresholds for species-specific enrichments.

**Figure 2:**
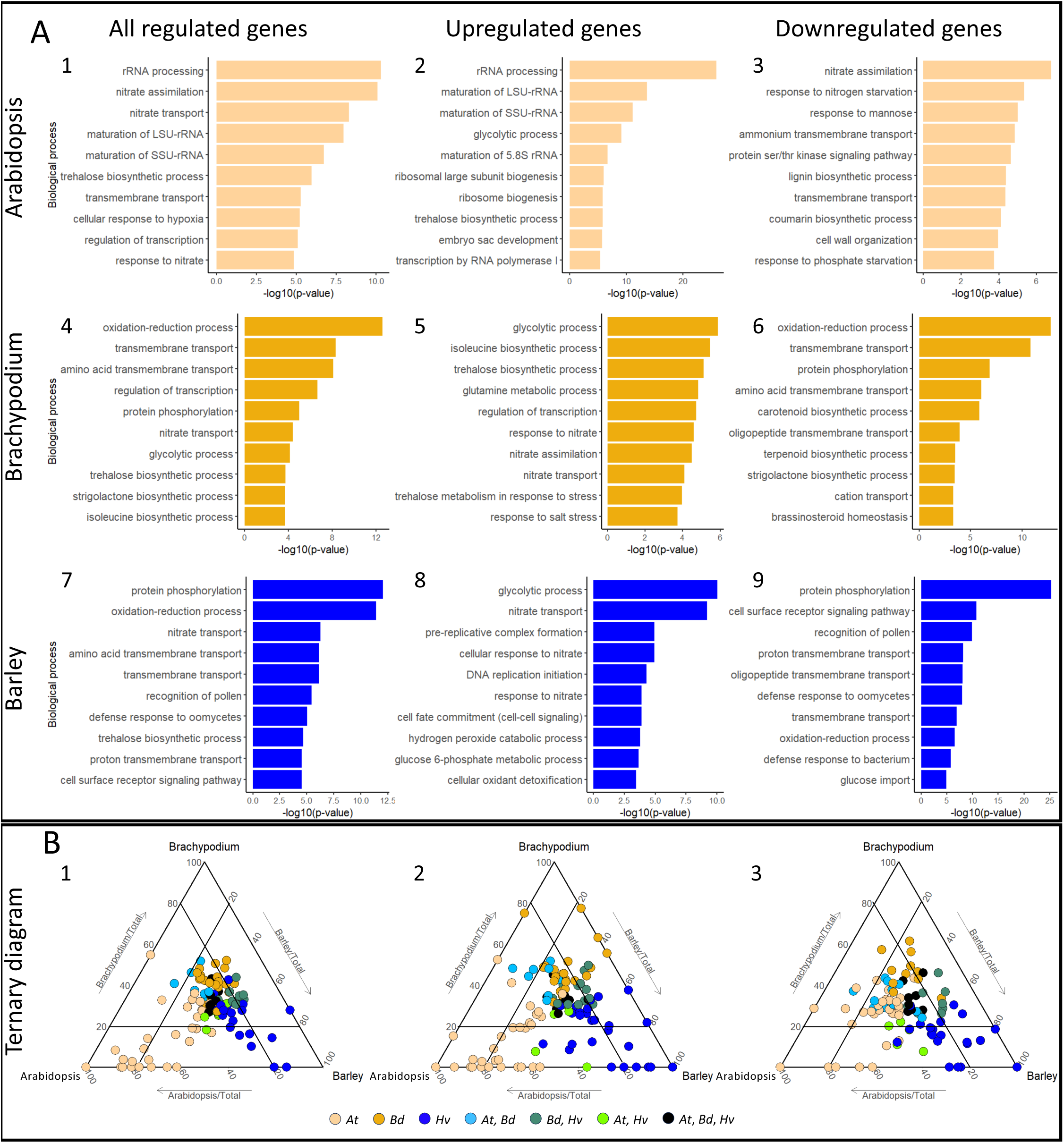
Gene Ontology (GO) enrichment analysis for Arabidopsis, Brachypodium and Barley after 1mM NO_3_ ^-^ treatment (combination of 1.5h and 3h). **(A)** The 10 best enriched GO terms (based on their p-value score) were plotted for each species. The full lists of enriched GO terms are available in Supp Table 3A. **(B)** Ternary diagrams for comparative enrichment analysis. Diagrams represent the distribution of GOs that belong to the 50 most-significantly enriched GO terms in at least one species (total of 109, 118 and 95 GO terms for B1, B2 and B3, respectively). Each axes represent the Normalized Enrichment Score (NES) for one species and each dot represents a GO term. Colors indicate for which species the GO term was significantly enriched in initial per-species analysis (ElimFisher test, pval<0.05). Internal bars represent a 20% threshold used for comparing enrichment scores between species (see Results). The lists of selected GOs and their NES are available in Supp Table 4A. For both (A) and (B), the first, second and third columns represents biological process GO terms considering all the DEGs, only the upregulated DEGs and only downregulated DEGs, respectively. Gene ontology terms, their enrichment scores and their p-values were generated using topGO R package.

As expected, N nutrition processes were highly enriched across species (“response to nitrate” [GO:0010167], “nitrate transport” [GO:0015706], “ammonium transmembrane transport” [GO:0072488] and “amino acid transmembrane transport” [GO:0003333]; Figure 2A). Carbon-related terms (“glycolytic process” [GO:0006096] and “trehalose biosynthetic process” [GO:0005992]) were also commonly enriched. Hormone pathways (“response to gibberellin” [GO:0009739] and “response to auxin” [GO:0009733]) were significantly enriched in all species, despite lower ranking (Supp Table 3A). On the other hand, some processes were enriched only in 1 or 2 species: translation-related (“rRNA processing” [GO:0006364], “maturation of SSU-rRNA” [GO:0000462], “maturation of LSU-rRNA” [GO:0000470], “ribosome biogenesis” [GO:0042254]) in Arabidopsis only (see below); “strigolactone biosynthetic process” [GO:1901601] in Brachypodium; “cell surface receptor signaling pathway” [GO:0007166] in barley; …

### Comparative analysis of regulated biological processes

Comparing GO enrichments required quantifying differences between species, as TopGO provided per-species enrichment but not inter-species comparison. A difficulty was due to size differences of GO databases between species: 85% of Arabidopsis genes were associated with at least one GO term, against 48% in Brachypodium and barley. For example, “nitrate assimilation” [GO:0042128] had 35, 6 and 3 detected genes in Arabidopsis, Brachypodium and barley, respectively (including 25, 5 and 2 DEGs); it was enriched significantly in Arabidopsis and Brachypodium only, due to low barley annotation (Figure 2A; Supp Table 3A). Similarly, “Abscisic acid-activated signaling pathway” [GO:0009738] was statistically enriched only in Arabidopsis, despite similar enrichment scores between the species (number of DEGs in GO term / number of detected genes in GO term; 71/243=0.29, 19/73=0.26 and 10/38=0.26 for Arabidopsis, Brachypodium and barley, respectively; Supp Table 4A1).

A pipeline for comparative analysis was developed based on TopGO per-species enrichment analyses and enrichment score comparisons between species. GO terms that 1/ contained at least 5 detected genes per species and 2/ belonged to the 50 most-significantly enriched terms in at least one species (TopGO p-values) were selected. Their Normalized Enrichment Scores (NES; see Materials and Methods) were calculated for each species and plotted on ternary diagrams (Figure 1). A threshold was arbitrarily defined (internal bars on Figure 1): GO terms with NES <20% were under-represented in a species; those >20% across all three species were non-specific (center of the diagram).

As for per-species enrichment analysis, comparative analysis used 1.5h and 3h DEGs for all DEGs, upregulated only or downregulated only (Figure 2B; Supp Table 4A), plus separate 1.5h and 3h analyses (Supp Figures 4 & 5; Supp Table 4B & 4C).

About half of GO terms clustered at the center of ternary diagrams (69/109, 54/118 and 49/95 for all DEGs, upregulated only and downregulated only, respectively; Figure 2B and Supp Table 4A), suggesting many nitrate-responsive biological processes were common across the three species. These included previously identified nitrate pathways (“nitrate transport” [GO:0015706], “response to nitrate” [GO:0010167] and “cellular response to nitrate” [GO:0071249]). Carbon pathways were also well represented (“glycolytic process” [GO:0006096], “pentose-phosphate shunt” [GO:0006098], “tricarboxylic acid cycle” [GO:0006099; in upregulated set] and “trehalose biosynthetic process” [GO:0005992]), confirming core nitrate response conservation. Phytohormone responses (cytokinin, auxin, gibberellin, jasmonic acid, karrikin and brassinosteroid) were also commonly regulated. Although “cytokinin metabolic process” [GO:0009690] and “cytokinin biosynthetic process” [GO:0009691] were common across species, “response to cytokinin” [GO:0009735] was specific to Arabidopsis and Brachypodium. Some specificities thus exist despite global conservation of nitrate effects on phytohormone pathways. Arabidopsis/Brachypodium similarities may relate to their undomesticated nature, contrary to barley.

In agreement with the GO per-species analysis, translation-linked processes were more enriched in Arabidopsis than in others: “maturation of 5.8S rRNA” [GO:0000460], “maturation of LSU-rRNA from tricistronic rRNA transcript (SSU-rRNA, 5.8S rRNA, LSU-rRNA)” [GO:0000463], “maturation of 5.8S rRNA from tricistronic rRNA transcript (SSU-rRNA, 5.8S rRNA, LSU-rRNA)” [GO:0000466], “maturation of LSU-rRNA” [GO:0000470], “transcription by RNA polymerase I” [GO:0006360] and “rRNA processing” [GO:0006364] represented 6 of the 8 Arabidopsis-specific GO terms (Supp Table 4A1). Arabidopsis and barley shared some additional processes related to translation: “maturation of SSU-rRNA from tricistronic rRNA transcript (SSU-rRNA, 5.8S rRNA, LSU-rRNA)” [GO:0000462], “maturation of SSU-rRNA” [GO:0030490] and “ribosomal large subunit biogenesis” [GO:0042273]. Similar results occurred with upregulated DEGs only (Supp Table 4A2). This suggests a stronger stimulation of translation in Arabidopsis during the timeframe of the experiment.

“Photosynthesis, light harvesting in photosystem I” [GO:0009768] responded specifically in Brachypodium/barley (no regulated gene out of 20 in Arabidopsis; Supp Table 4A1), as did “photomorphogenesis” [GO:0009640] to a lesser extent. Amino acid biosynthesis processes (“cysteine biosynthetic process from serine” [GO:0006535]; “lysine biosynthetic process via diaminopimelate” [GO:0009089]) and cell wall (“glucuronoxylan biosynthetic process” [GO:0010417]) also showed similar patterns.

Upregulated DEGs analysis (Supp Table 4A2) showed that most nitrate-related GO terms were stimulated across all three species, apart from “nitrate transport” [GO:0015706]. This term was more enriched in Brachypodium/barley only, possibly linked to the high NRT transporter divergence between Arabidopsis and cereals (Plett *et al*., 2010).

Downregulated DEGs analysis (Supp Table 4A3) identified “response to starvation” [GO:0042594], “cellular response to nitrogen starvation” [GO:0006995] and “cellular response to phosphate starvation” [GO:0016036] as much more enriched in Arabidopsis/barley. This suggests a stronger activation of starvation-processes during starvation pre-treatment, and could reflect the adaptation of these species to rich soil, whereas Brachypodium is adapted to dry and poor soils (Des Marais and Juenger, 2015). Further gene-level analysis of these pathways could clarify the role of this repression.

### Generation of orthogroups towards a comparative analysis of regulated genes

To further investigate conservation and divergence of responses at the gene level, it was necessary to compare regulation between orthologs from the three species. We performed a whole-genome analysis using Orthofinder (Emms and Kelly, 2015) to cluster orthologs identified by protein sequence similarity into orthogroups (OGps). A total of 36 754 OGps were identified (Supp Table 5), including 9 974 OGps composed of genes from the three species (27% of OGps, regrouping 43% of genes), 6 509 OGps encompassing two species (18% and 21%), 4 741 OGps composed of paralogs from a single species (13% and 21%) and 15 530 OGps composed of a single gene (42% and 15%) (Supp Figure 6). As expected from phylogenetic relationships, more genes were grouped into Brachypodium/barley bi-species OGps than into Arabidopsis-containing bi-species OGps.

DEGs tended to be over-represented in tri-species OGps and under-represented in mono-species OGps compared to the distribution of whole-genome gene models (Supp Figure 6), suggesting that nitrate-regulated sets are enriched in conserved genes. Nitrate-responsive genes were also over-represented in Brachypodium/barley bi-species OGps, suggesting that part of the nitrate response in these species is carried by specific genes divergent from Arabidopsis. Biological functions of DEGs belonging to Brachypodium/barley bi-species OGps were highly enriched in GO terms “protein phosphorylation” [GO:0006468] and “regulation of transcription, DNA-templated” [GO:0006355] (Supp Figure 7 and Supp Table 6). These genes encode kinases (MAPKs, WAKs, LRRs, S6Ks, …) and transcription factors (NACs, bHLHs, bZIPs, WRKYs, …) that could be involved in nitrate response specifically in Pooideae. However, they are not ideal candidates for reverse-genetic approaches, as 1/ their list still contains hundreds of genes (for instance, more than 150 Brachypodium transcription factors) and 2/ the method cannot exclude that a fairly divergent ortholog carries a similar function in Arabidopsis. Studying divergent regulation within tri-species OGps is therefore a better approach to identify Pooideae-specific nitrate responses (see below).

### Identification of conserved and divergent responses of known genes involved in nitrate-related pathways

Gene regulation was analyzed in OGps containing known Arabidopsis nitrate-responsive genes. The 50 most consistently nitrate-regulated genes, identified from a meta-analysis of 27 transcriptomic datasets (Canales *et al*., 2014), were investigated (Table 1). The 50 genes belonged to 41 OGps, grouping 89 Arabidopsis genes; 70 of these were regulated in our experiment. In particular, 49 of the initial 50 genes were induced, which is a good indicator of the quality of our Arabidopsis transcriptomic analysis. Orthologs in Brachypodium and barley were largely regulated as well: 86% of tri-species OGps (19/28) responded in at least two species (Table 1), suggesting high conservation of core nitrate-responsive genes in plants. The NAS OGp (encoding a key enzyme in nicotianamine biosynthesis) was of particular interest: while 3 of 4 paralogs were induced in Arabidopsis, none of the 3 Brachypodium and 12 barley orthologs were regulated (Table 1). Nicotianamine is necessary for transport of Fe^2+^ and other cations to maintain metal homeostasis in plants. Since Arabidopsis and grasses have different strategies to acquire Fe from soil (Marschner and Römheld, 1994), our results suggest they also differ in maintaining metal homeostasis in response to nitrate. The much higher number of paralogs in barley compared to the other two species could reflect domestication effects on this gene family, likely independent of nitrate response itself.

**Table 1:**
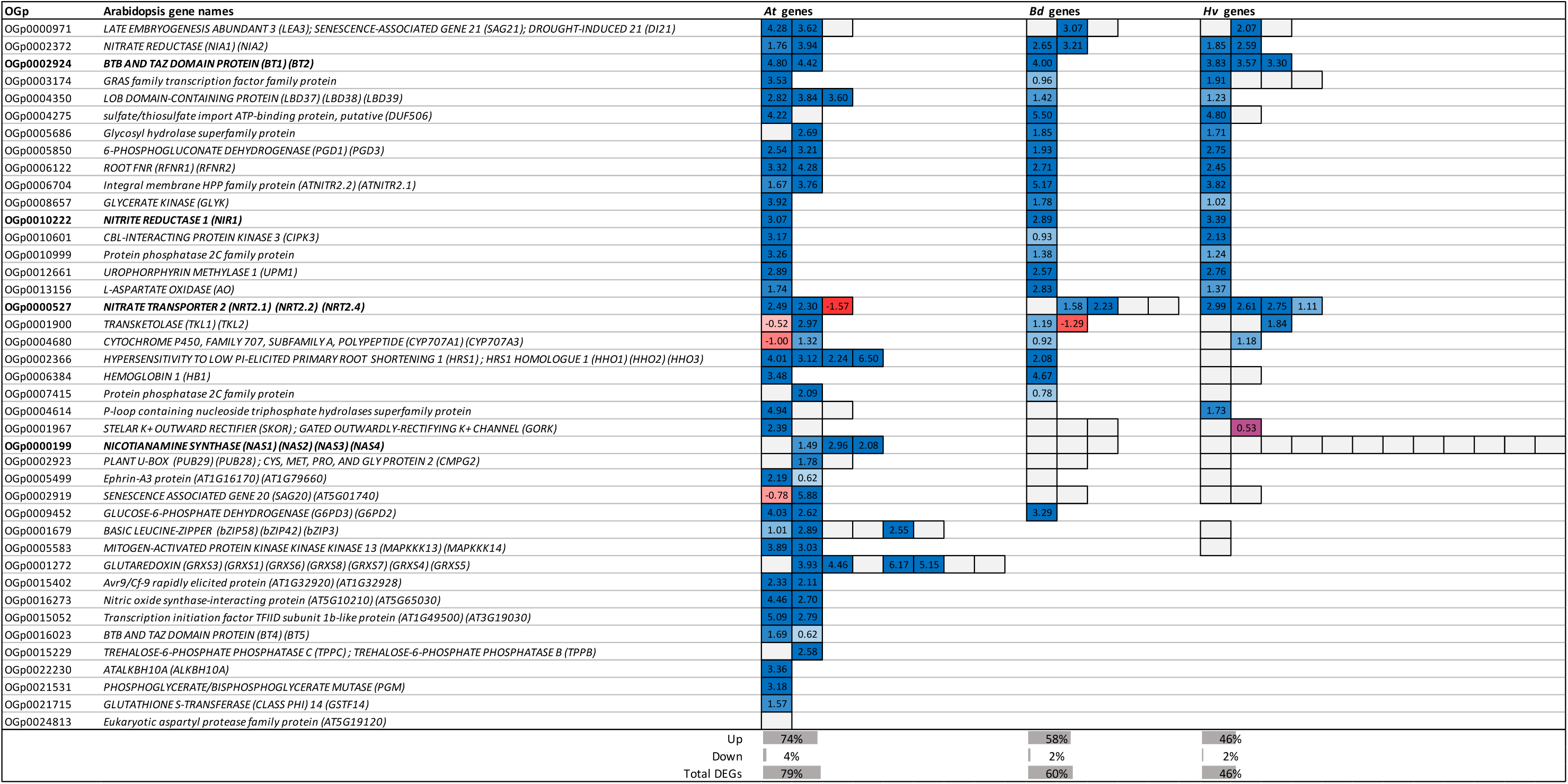
Nitrate-response of top 50 most consistent genes regulated by nitrate in Arabidopsis (list from Canales et al., 2014) and of their orthologs in Brachypodium and barley. Lines represent OGps; colored rectangles represent genes, grouped by species (*At*, Arabidopsis; *Bd*, Brachypodium; *Hv*, barley). Color scale and numbers indicate maximum Log_2_FC values (1.5h or 3h nitrate *vs* mock); blue, upregulated; red, downregulated; purple, upregulated at 1.5h / downregulated at 3h; grey, not-regulated/not-detected. OGps in bold are discussed in the text. Proportions of upregulated, downregulated and total DEGs among represented genes are indicated below the table for each species.

As a complementary approach, regulation of main genes involved in nitrate transport, assimilation and signaling (Vidal et al., 2020) was investigated (Table 2, organized into 12 physiological processes). Among “**transport and assimilation**” processes, most OGps were similarly regulated in the three species (Table 2). As expected, *AtNIA1*, *AtNIA2* and *AtNIR*, encoding enzymes required for nitrate assimilation, were upregulated. Two of the three AtNRT1.1 orthologs in each Pooideae species were upregulated, suggesting high capacity of response for nitrate uptake. Genes of the *AtNRT2.1* orthogroup, encoding the main high-affinity nitrate transporter, were also regulated in all three species. Interestingly, all barley genes of this OGp were upregulated, again suggesting a strong nitrate uptake response. This is consistent with previous observations of higher response of high affinity nitrate uptake in barley than in Brachypodium (David *et al*., 2019). The *AtNRT2.1* OGp also contains *AtNRT2.4*, involved in nitrate uptake specifically under severe nitrate limitation (Kiba *et al*., 2012), which was downregulated as expected. The absence of downregulation of Brachypodium or barley genes in this OGp suggests no ortholog shares this specific function.

**Table 2:**
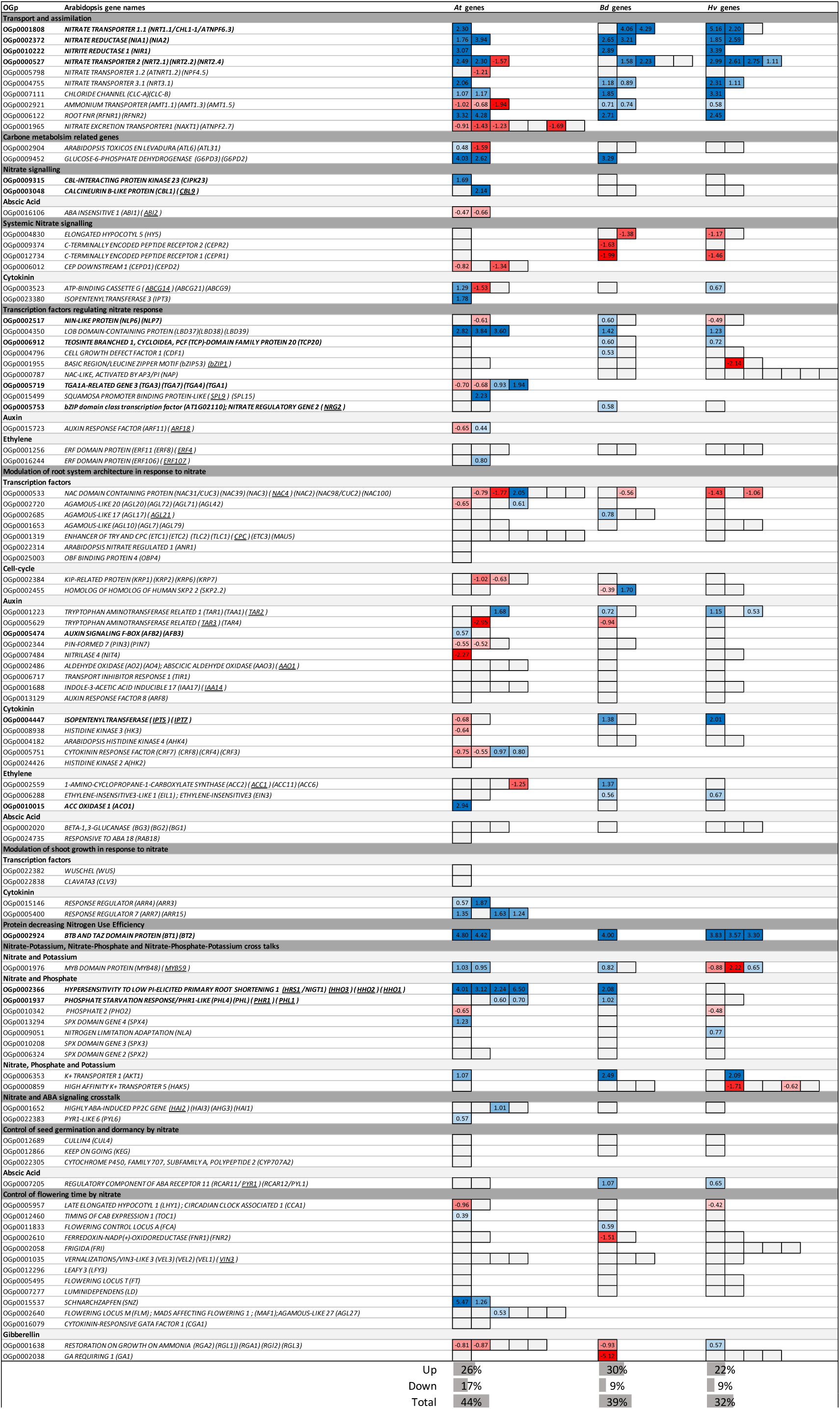
Nitrate-response of genes involved in nitrate transport, assimilation and signaling pathway (list from Vidal *et al*., 2020). Lines represent OGps, organized by physiological processes; colored rectangles represent genes, grouped by species (*At*, Arabidopsis; *Bd*, Brachypodium; *Hv*, barley). Color scale and numbers indicate maximum Log_2_FC values (1.5h or 3h nitrate *vs* mock); blue, upregulated; red, downregulated; grey, not-regulated/not-detected. OGps in bold are discussed in the text; underlined genes have been discussed in Vidal *et al*. (2020). Proportions of upregulated, downregulated and total DEGs among represented genes are indicated below the table for each species.

Genes coding for ammonium transporters were downregulated in response to nitrate in Arabidopsis (Table 2), whereas identified orthologs in Brachypodium and barley were upregulated. The three species are known to be ammonium-sensitive, compared to ammonium-insensitive species such as rice (Britto *et al*., 2001; Loqué *et al*., 2006). In Arabidopsis, ammonium transporters are induced by N starvation (Loqué and von Wirén, 2004), consistent with repression upon nitrate supply to N-starved plants in our experiment. The divergent response may reflect physiological differences, with Pooideae being less sensitive to ammonium than Arabidopsis.

**Nitrate signaling** requires several actors, such as NRT1.1, calcium signals and kinases. Under low nitrate, AtCIPK23 and AtCBL9 interact and phosphorylate AtNRT1.1 to switch to high-affinity nitrate transport activity. *AtCIPK23* is induced by nitrate (Ho *et al*., 2009), whereas transcriptional regulation of *AtCBL9* by nitrate is not documented. In our experiment, both genes were upregulated in response to nitrate, but their closest orthologs in Brachypodium and barley were not (Table 2), suggesting at least partial divergence of nitrate-signaling mechanisms between species. This does not exclude a role for AtCIPK23 and AtCBL9 orthologs in nitrate response, as their function may be regulated post-transcriptionally.

Nitrate response mobilizes many **transcription factors**. Among them, AtNLP6 and AtNLP7 are major regulators (Marchive *et al*., 2013), and AtNLP7 also acts as an intracellular nitrate sensor (Liu *et al*., 2022). *AtNLP7* is described either as transcriptionally insensitive to N availability (Castaings *et al*., 2009) or transiently upregulated by nitrate, peaking 30 min after addition (Varala *et al*., 2018). In our conditions, *AtNLP7* and *HvNLP7* were slightly downregulated, while *BdNLP7* was slightly upregulated (Table 2). AtTCP20 transcription factor interacts with AtNLP6 and AtNLP7 and controls nitrate response of cell cycle and root growth (Guan *et al*., 2014; Guan *et al*., 2017). To our knowledge, no study has shown transcriptional regulation of *AtTCP20* by nitrate in Arabidopsis or Pooideae. In our data *AtTCP20* was not regulated in Arabidopsis, but two Pooideae orthologs were upregulated (Table 2). *AtNRG2* encodes another transcription factor interacting with AtNLP7 and mediating nitrate signaling; its expression is not known to be regulated by nitrate, ammonium or N deprivation (Xu *et al*., 2016). Interestingly, the unique *AtNRG2* ortholog in Brachypodium was regulated in our experiment (Table 2). Together, these observations suggest an additional level of transcriptional regulation in Pooideae compared to Arabidopsis: TCP20 and NRG2 likely act downstream of another transcriptional regulator in Pooideae.

TGA transcription factors, especially AtTGA1 and AtTGA4, play important roles in Arabidopsis nitrate response: they bind *AtNRT2.1* and *AtNRT2.2* promoters and regulate lateral root elongation (Alvarez *et al*., 2014). Surprisingly, no TGA orthologs were identified by our OGp method in Brachypodium or barley (Table 2). However, BLAST analyses identified numerous *TGA* genes in Brachypodium and barley, several being regulated by nitrate similarly to *AtTGA1-4* (*e.g.* Bradi1g46060 [Log_2_FC=-1.45], Bradi3g15590 [-0.74], Bradi3g60870 [+1.12], Bradi2g23890 [+2.73], HORVU4Hr1G025130 [-0.79], HORVU5Hr1G053330 [-0.66], HORVU3Hr1G084340 [+1.08] and HORVU1Hr1G073870 [+3.56]). These *TGA* orthologs were classified in mono-species or bi-species Bd+Hv OGps. This illustrates both strength and weakness of the OGp approach: it groups close orthologs by protein similarity but can exclude moderate divergent genes. For *TGA*s, protein sequences are highly divergent between closest orthologs (around 50% identity), but functions are probably conserved, as they respond similarly to nitrate. This reasoning of divergent orthologs with similar function may apply to all mono-species OGps, such as those containing *AtSPL9*, *AtANR1*, *AtCRF* and other hormone-pathway regulators (*ARFs*, *ABI*, …; Table 2). Alternatively, these genes could be truly Arabidopsis-specific, lacking both close orthologs in other species and distant ortholog with similar nitrate-response function. Detailed and exhaustive analyses of candidate gene families would be required to distinguish between these possibilities.

Auxin, cytokinin, ethylene and abscisic acid (ABA) pathways contribute to nitrate regulation of **root system architecture**, a complex trait differing between monocots and dicots (Hochholdinger *et al*., 2004; Osmont *et al*., 2007). *AtAFB3* encodes an auxin receptor transcriptionally regulated by nitrate and controlling root system architecture in Arabidopsis (Vidal *et al*., 2010). In our conditions, *AtAFB3* was upregulated, but orthologs in Brachypodium and barley were not nitrate-responsive (Table 2), suggesting divergence in nitrate-dependent regulation of root system architecture between Arabidopsis and Pooideae. This is in agreement with reported differences between Arabidopsis and Brachypodium in the control of root architecture by auxin, brassinosteroids and ethylene (Hardtke and Villalobos, 2015). Further analysis of auxin pathways in nitrate response in Pooideae would help clarify the role of AtAFB3 orthologs.

A role of ethylene in root development in response to nitrate has been identified in Arabidopsis: nitrate induces *AtACO1*, encoding an ethylene biosynthesis enzyme (Tian *et al*., 2009), leading to rapid ethylene increase in lateral root primordia and repression of lateral roots under high nitrate. In our analysis, *AtACO1* expression was induced by nitrate, whereas Brachypodium and barley orthologs were insensitive (Table 2), again suggesting different mechanisms between Arabidopsis and Pooideae. Cytokinins, beyond systemic nitrate signaling, are involved in primary root development in response to nitrate (Naulin *et al*., 2020) *via* the regulation of *IPT* genes required for cytokinin biosynthesis. In particular, *AtIPT5* is regulated by nitrate addition (Miyawaki *et al*., 2004). In our experiment, *AtIPT5* was downregulated after 3h (Table 2), likely reflecting repression by products of nitrate assimilation. Brachypodium and barley orthologs were upregulated by nitrate. This may again indicate divergence in regulation of root architecture, or a slower establishment of negative feedback in Pooideae compared to Arabidopsis.

BT2, a member of the Bric-a-Brac/Tramtrack/Broad family of scaffold proteins known to be essential in female and male gametophyte development (Gingerich *et al*., 2007; Robert *et al*., 2009), has also been described as regulating nitrate uptake and **decreasing N use efficiency** (Araus *et al*., 2016). In our experiment, all Arabidopsis, Brachypodium and barley members of this orthogroup were highly induced by nitrate (Tables 1 & 2).

### Identification of conserved and divergent orthologous gene responses based on comparative analysis of regulated biological processes

To identify specificities and generalities between species specifically linked to gene regulation – and not to presence/absence of orthologs – the analysis of transcriptomic response was restricted to tri-species OGps. Gene ontology (GO) and orthology (OGp) approaches were then combined to examine DEGs. We used a similar pipeline as for the comparative analysis of whole DEG sets (Figure 1), restricting input DEG lists and Ensembl databases to genes belonging to tri-species OGps. Ternary diagrams of Normalized Enrichment Scores were obtained for 1/ all DEGs, 2/ only upregulated DEGs and 3/ only downregulated genes (Supp Figure 8 and Supp Table 7). Forty to sixty percent of GO terms were equally enriched in the three species, clustering at the center of the diagrams, while the remaining GO terms were more enriched in one or two species. Interestingly, all pairs of species were associated with enrichment in some biological processes.

Based on these diagrams, we selected a subset of GO terms for detailed gene-level analysis using our orthology data. This approach corrects the drawback of unequal GO annotation quality between species, as genes of the same orthogroup are considered independently of their annotation.

#### Response of N-related processes is mainly conserved between species

In agreement with the comparative analysis based on whole DEG sets, the GO terms “response to nitrate” [GO:0010167], “nitrate transport” [GO:0015706] and “cellular response to nitrate” [GO:0071249] were commonly enriched between species when restricting the analysis to tri-species OGps (Supp Table 7). Examining regulation of orthologs belonging to these GO terms and tri-species OGps confirmed that nitrate response was well conserved at the gene level (Supp Table 8). Most of these genes are included in the list published by Vidal et al. (2020) (see above).

According to the comparative analysis based on tri-species OGps, the “**cellular response to nitrogen starvation**” [GO:0006995] process was specific to Arabidopsis and barley (Supp Table 7). However, when considering the annotated genes in this category and their orthologs, this GO term appeared downregulated in all three species (Supp Table 9A). This illustrates the importance of combining GO and orthology analyses to avoid biases due to unequal GO annotation between species.

The GO term “**amino acid transmembrane transport**” [GO:0003333] was enriched in all species (Supp Table 7). Corresponding genes and orthologs were mainly downregulated, and the proportion of regulated genes was similar between species (Supp Table 9B). However, only 20% of regulated OGps (5/25) were regulated in all three species, whereas 52% (13/25) and 28% (7/25) responded in only one and two species, respectively. These specifically responsive OGps include strongly regulated genes: the closest ortholog of AtANT1, known in Arabidopsis to transport aromatic amino acids, neutral amino acids, arginine and auxin (Chen *et al*., 2001), is induced only in Brachypodium; a non-characterized amino acid transporter (AT3G54830) is highly and specifically upregulated in Arabidopsis. Thus, amino acid transporters might include key actors of nitrate response in a species-specific manner.

#### Regulation of gibberellin biosynthesis pathway is common to Brachypodium and barley

Gibberellins (GA) have been implicated in the control of root architecture by nitrate supply (Camut *et al*., 2021). The “**gibberellin biosynthetic process**” GO term [GO:0009686] was specifically enriched in Brachypodium (Supp Table 7A) but also appeared regulated in barley when analyzing orthologs (Table 3). Most regulated genes were repressed by nitrate (10/11 and 5/8 in Brachypodium and barley, respectively). In contrast, “response to gibberellin (GA)” [GO:0009739] was common to all three species (Supp Table 7A), which was confirmed at gene level using tri-species OGps (Supp Table 10). This suggests that despite a common GA response downstream of nitrate treatment, GA biosynthesis is differentially regulated in Pooideae compared to Arabidopsis.

**Table 3:**
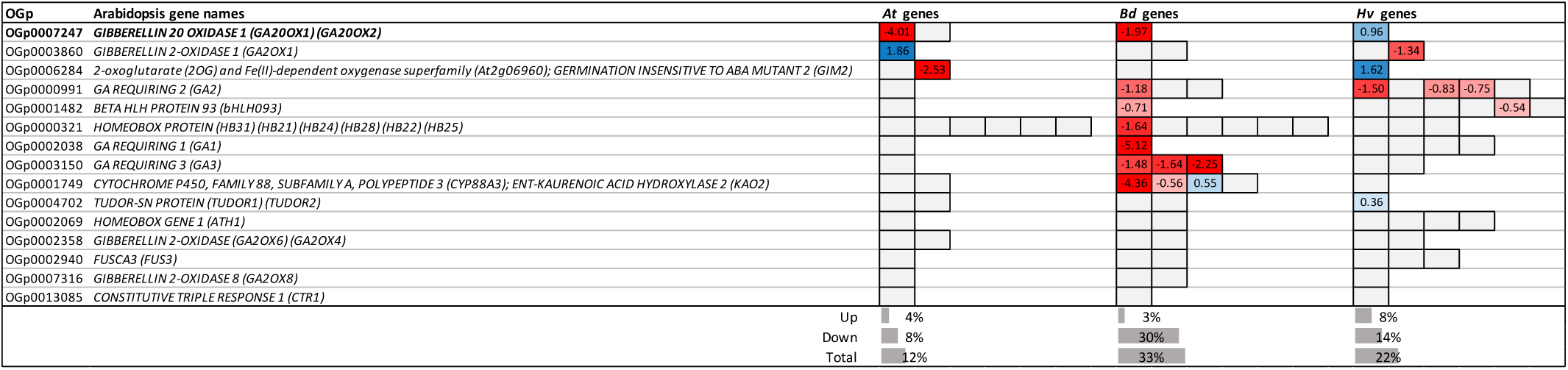
Gene nitrate-response in tri-species OGps belonging to “gibberellin biosynthetic process” GO term [GO:0009686]. The GO term was identified as Brachypodium-specific by the comparative analysis on tri-species OGps. Lines represent OGps; colored rectangles represent genes, grouped by species (*At*, Arabidopsis; *Bd*, Brachypodium; *Hv*, barley). Color scale and numbers indicate maximum Log_2_FC values (1.5h or 3h nitrate *vs* mock); blue, upregulated; red, downregulated; grey, not-regulated/not-detected. OGps in bold are discussed in the text. Proportions of upregulated, downregulated and total DEGs among represented genes are indicated below the table for each species.

The case of *GA20OX1*, encoding an enzyme involved in the last steps of gibberellin biosynthesis (Rieu *et al*., 2008), was peculiar: *AtGA20OX1* and its closest ortholog in Brachypodium were highly repressed by nitrate, whereas the ortholog in barley was upregulated (Table 3). This gene is known to be a key selection locus of the Green Revolution (Jia *et al*., 2009): modern domesticated plants such as rice, barley and wheat carry mutations leading to semi-dwarf phenotypes selected to avoid lodging. Thus, divergent nitrate response of this gene between domesticated and non-domesticated species could be linked to domestication.

#### Nitrate and phosphate crosstalk

Recent studies in Arabidopsis, rice and wheat show that phosphate (P) starvation response occurs only in the presence of nitrate (Medici *et al*., 2015; Medici *et al*., 2019; Dissanayake *et al*., 2019; Hu *et al*., 2019). The GO term “**cellular response to phosphate starvation**” [GO:0016036] was more enriched in response to nitrate in Arabidopsis and barley than in Brachypodium (Supp Table 7). However, ortholog response analysis did not fully confirm this, as the process appears more enriched in Arabidopsis and evenly responsive in Brachypodium and barley (Table 4).

**Table 4:**
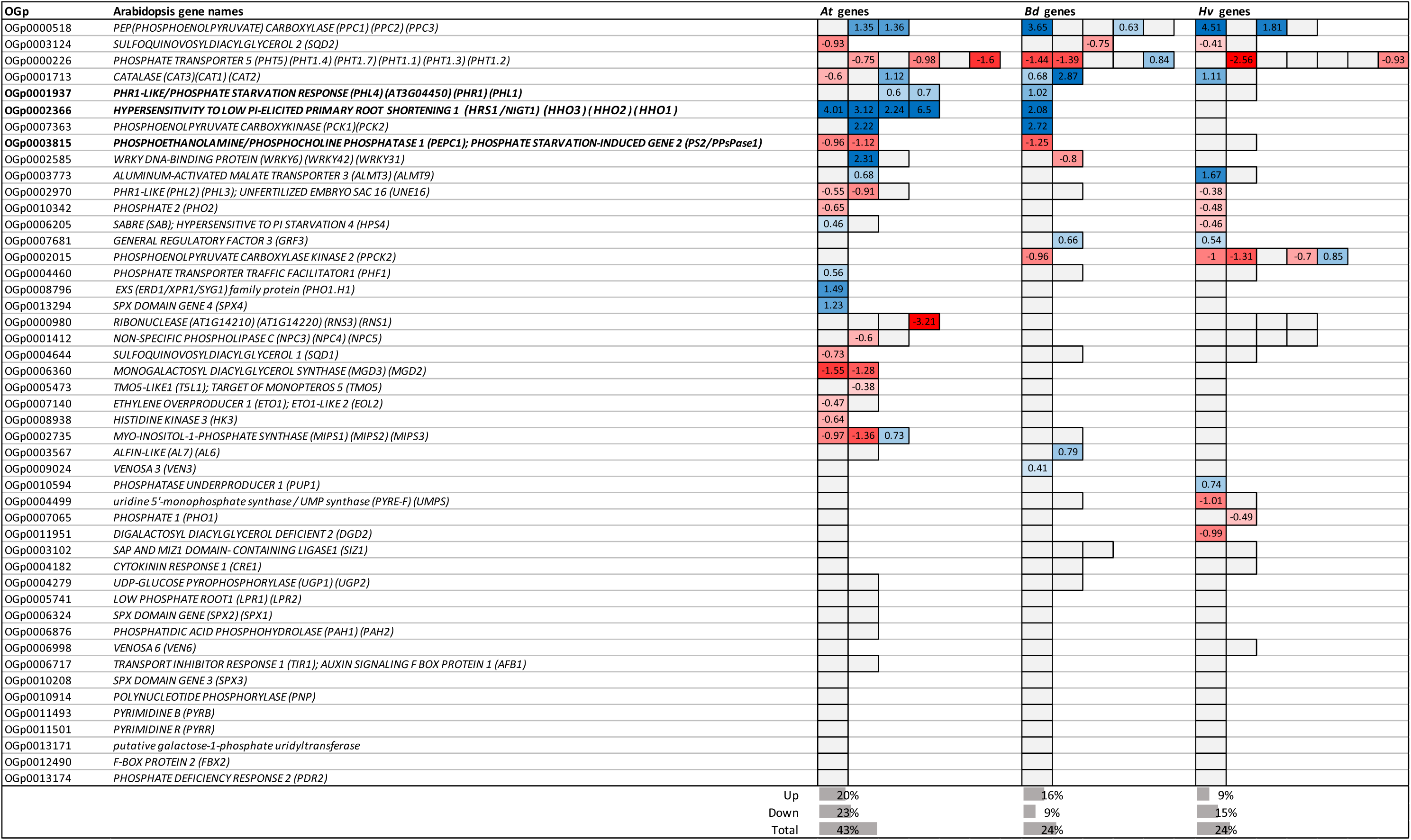
Gene nitrate-response in tri-species OGps belonging to “cellular response to phosphate starvation” GO term [GO:0016036]. The GO term was identified as common to Arabidopsis and barley by the comparative analysis on tri-species OGps. Genes associated to this GO term and belonging to tri-species OGps were extracted. Lines represent OGps; colored rectangles represent genes, grouped by species (*At*, Arabidopsis; *Bd*, Brachypodium; *Hv*, barley). Color scale and numbers indicate maximum Log_2_FC values (1.5h or 3h nitrate *vs* mock); blue, upregulated; red, downregulated; grey, not-regulated/not-detected. OGps in bold are discussed in the text. Proportions of upregulated, downregulated and total DEGs among represented genes are indicated below the table for each species.

Phosphate deficiency inhibit Arabidopsis primary root growth through the action of the transcription factors AtHRS1/NIGT1 and AtHHO1, but only in presence of nitrate (Medici et al., 2015). In turn, the nitrate-induction of *AtHRS1/NIGT1* is enhanced by phosphate deficiency, *via* activation of AtPHR1 transcription factor (Maeda et al., 2018). In our experiment, all four Arabidopsis genes and the unique Brachypodium gene of the *HRS1* orthogroup were upregulated, whereas the single barley gene was unresponsive (Tables 2 & 4). *AtPHR1*, one paralog in Arabidopsis (*AtPHL1*) and their unique ortholog in Brachypodium were slightly upregulated, but the barley ortholog was insensitive (Tables 2 & 4). Barley orthologs in these two OGps are thus not likely carrying the functions described in Arabidopsis. A similar situation has been observed in wheat, where the closest *AtHRS1/NIGT1* ortholog was regulated later than in Arabidopsis, while a more distant ortholog responded rapidly (Dissanayake et al., 2019). Such cases may indicate domestication-linked molecular divergence, with a function carried by relatively distant orthologs.

Arabidopsis *AtPEPC1* and *AtPPsPase1* – belonging to the same OGp – were both downregulated by nitrate, as was the closest Brachypodium ortholog, but not the two barley ones (Table 4). These Arabidopsis phosphatases participate in recycling lipid membrane polar heads triggered by P starvation (Hanchi *et al*., 2020) and, to our knowledge, have not been linked to N-responses. They therefore constitute candidates for new molecular players of N–P crosstalk in Arabidopsis and Brachypodium.

#### rRNA processing is activated in response to nitrate almost exclusively in Arabidopsis

The “**rRNA processing**” GO term [GO:0006364] appeared highly enriched specifically in Arabidopsis when restricting the analysis to tri-species OGps, consistent with per-species analysis and whole-DEG comparative analysis (see above). Comparison of orthologs from tri-species OGps confirmed that rRNA processing genes were largely induced in Arabidopsis in a very specific manner: 45% (108/242), 2% (4/248) and 7% (19/256) of genes were induced in Arabidopsis, Brachypodium and barley, respectively (Supp Table 10). None of the genes were repressed by nitrate. Since rRNAs are required for translation, this apparent Arabidopsis specificity may reflect timing differences, with Arabidopsis being at a more advanced response stage.

#### Genes related to membrane trafficking and cell wall biogenesis are differentially regulated in response to nitrate

The GO term “**Rab protein signal transduction**” [GO:0032482] was specifically enriched in Brachypodium in the comparative analysis of upregulated genes restricted to tri-species OGps (Supp Table 7B). Surprisingly, corresponding DEGs were mainly upregulated in Brachypodium (7/7) and downregulated in Arabidopsis (7/9), whereas barley orthologs responded weakly (1 up and 1 down) (Supp Table 12A). Notably, upregulated Arabidopsis genes and downregulated Brachypodium genes are not direct orthologs. These regulated genes mainly encode Rab GTPases of RabA, B and C subfamilies, involved in vesicle trafficking (Woollard and Moore, 2008). To our knowledge, no role of Rab proteins in nitrate response has been described. They are however important for abiotic stress responses (Tripathy et al., 2021) and for growth through deposition of enzymes and cell wall components (Lycett, 2008). Interestingly, some cell wall related processes are also mostly downregulated in Arabidopsis, while only few orthologs are regulated in Brachypodium and barley (Supp Tables 11B & 11C). This could indicate differences in early growth responses to nitrate between species, involving modification of cell wall properties.

#### Identification of pathways enriched exclusively in Brachypodium and barley

The GO term “**cysteine biosynthetic process from serine**” [GO:0006535] was specifically enriched in Brachypodium and barley (Supp Table 7), as confirmed by the heatmap of tri-species OGps (Table 5A). In particular, two enzymes in this pathway were specifically upregulated in Pooideae: serine acetyltransferase (SAT) and O-acetyl(thiol)lyase (OASTL), suggesting higher cysteine production in response to nitrate in Pooideae. OASTL requires pyridoxal phosphate (PLP) as cofactor (Hesse *et al*., 1999). Very interestingly, “**pyridoxal phosphate biosynthetic process**” [GO:0042823] was also specifically enriched in Brachypodium and barley, when considering upregulated DEGs (Supp Table 7B), which was confirmed at gene level (Table 5B). As cysteine is a product of sulfur assimilation, we checked sulfur-related GO terms. “Sulfur compound metabolic process” [GO:0006790] appeared evenly enriched in the three species (Supp Table 7), suggesting no imbalance in sulfur availability.

**Table 5:**
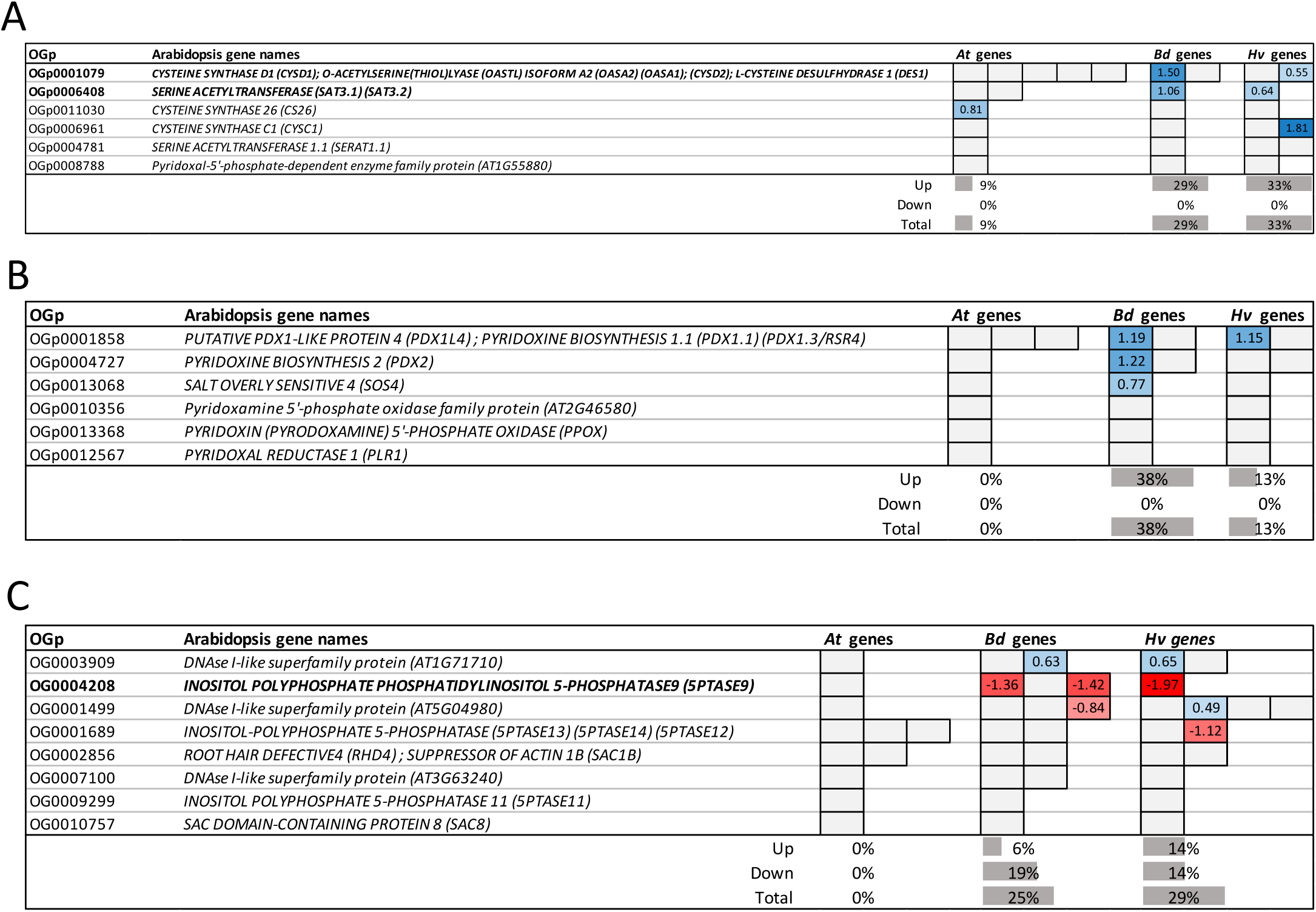
Gene nitrate-response in selected tri-species OGps from Pooideae-specific GO terms. GO terms “cysteine biosynthetic process from serine” [GO:0006535] **(A)** “pyridoxal phosphate biosynthetic process” [GO:0042823] **(B)** and “phosphatidylinositol dephosphorylation” [GO:0046856] **(C)** were identified by the comparative analysis on tri-species OGps as common to the two *Pooideae* species. Genes associated to those GO terms and belonging to tri-species OGps were extracted. Lines represent OGps; colored rectangles represent genes, grouped by species (*At*, Arabidopsis; *Bd*, Brachypodium; *Hv*, barley). Color scale and numbers indicate maximum Log_2_FC values (1.5h or 3h nitrate *vs* mock); blue, upregulated; red, downregulated; grey, not-regulated/not-detected. OGps in bold are discussed in the text. Proportions of upregulated, downregulated and total DEGs among represented genes are indicated below the tables for each species.

The GO term “**phosphatidylinositol dephosphorylation**” [GO:0046856] was also specifically regulated in Brachypodium and barley: no gene of the corresponding OGps was regulated in Arabidopsis, whereas 25% and 29% were responsive in the Pooideae species (Table 5C). Phosphorylated forms of phosphatidylinositol play important roles in lipid signaling, cell signaling and membrane trafficking. The large *5PTASE* family encodes phosphatases that hydrolyze 5-phosphates from phosphatidylinositol phosphates. Orthologs of *At5PTASE9* were downregulated in response to nitrate in Brachypodium and barley (Table 5C). In Arabidopsis, At5PTASE9 is involved in salt tolerance: *At5ptase9* mutants show decreased calcium influx and reduced ROS production, leading to lack of induction of salt-responsive genes and increased salt tolerance (Golani *et al*., 2013). Orthologs of At5PTASE9 in Brachypodium and barley could therefore regulate calcium fluxes and ROS production, both known to participate in nitrate signaling (Chu et al., 2021; Mao et al., 2025; Riveras et al., 2015).

## Discussion

The aim of this study was to compare the transcriptional nitrate response of Brachypodium and barley, as Pooideae members, to the model species Arabidopsis. We performed RNAseq on root tissues of plantlets after 1.5h and 3h of nitrate treatment to capture the most exhaustive transcriptional response. Our first analysis consisted of describing DEGs using gene ontology enrichment. Per-species analysis (Figure 2A) validated the experimental setup, since GO terms related to nitrate response were enriched in all three species. As expected, GO terms related to carbon metabolism or hormone signaling were also enriched. These observations were confirmed by the comparative analysis of whole DEG sets (Figure 2B), which identified these GO terms as commonly regulated across species. Comparative analysis also provided initial clues to identify GO terms enriched in only one species (*ie*, “rRNA processing” [GO:0006364], specifically enriched in Arabidopsis).

To deepen the analysis at the gene level, OGps were constituted to identify orthologous genes in the three species. DEGs were over-represented in bi- and tri-species OGps (Supp Figure 6), indicating that nitrate response involves predominantly conserved genes. DEGs belonging to Brachypodium/barley bi-species OGps were enriched in protein phosphorylation and regulation of transcription processes (Supp Figure 7), suggesting that some nitrate signaling actors such as kinases and transcription factors are specific to Pooideae.

We subsequently investigated OGps corresponding to known Arabidopsis genes responding to nitrate treatment (Table 1) or involved in nitrate metabolism and/or response (Table 2). Within these lists, respectively 57% and 30% of OGps showed a similar regulation pattern across the three species. Nitrate response seems globally well conserved between Arabidopsis and Pooideae based on core genes such as enzymes required for nitrate transport and assimilation. Some differences in regulation were noted, suggesting partial divergence of mechanisms between species. This supports the importance of combining crops and model plants to identify candidates that could enhance nitrate response. Some divergences are supported by known physiological differences between Arabidopsis and Pooideae, such as differentially regulated *NAS* genes involved in Fe homeostasis in response to nitrate, supporting a different strategy of Fe acquisition between Arabidopsis and grasses (Marschner and Römheld, 1994). Similarly, differences in gene regulation in hormone pathways necessary for root system architecture support different root phenotypes in response to nitrate between Arabidopsis and cereals (Smith and de Smet, 2012; Shahzad and Amtmann, 2017).

The last part of the study aimed to identify specific/common pathways and genes by focusing on conserved genes (tri-species OGps), thus reducing biases linked to divergence in protein sequences. The approach highlighted response divergence in cysteine biosynthesis from serine, which was more activated in Brachypodium and barley (Table 5A). This activation was further supported by specific induction of genes necessary for PLP biosynthesis, required for OASTL function (an enzyme required for cysteine biosynthesis), in Brachypodium and barley. We thus identify an intriguing Pooideae-nitrate-specific process that can lead to a better understanding of nitrate response in cereals.

Differences in how barley responds to nitrate compared to Brachypodium and Arabidopsis might be linked to domestication, which could have reshaped specific molecular and physiological traits, particularly in pathways critical for nutrient use and growth regulation. For instance, the gene *GA20OX1*, which plays a central role in gibberellin biosynthesis and contributes to the semi-dwarf phenotype selected during the Green Revolution, exhibits opposite regulation by nitrate in barley (upregulated) versus wild species (repressed). This divergence suggests that breeding for agronomic traits like reduced plant height has indirectly altered how barley manages N resources. Additionally, barley’s ortholog of AtHRS1 – a transcription factor involved in phosphate-nitrate crosstalk – is unresponsive to nitrate, unlike those in Brachypodium and Arabidopsis. This points to domestication-driven changes in how barley integrates N and P signaling, possibly reflecting adaptations to high-input agricultural environments. Furthermore, differences in regulation of N starvation responses and root architecture-related hormones (e.g., gibberellins and auxins) between barley and wild species underscore how domestication has fine-tuned metabolic and developmental pathways to optimize yield under cultivated conditions. These findings not only reveal the molecular footprint of domestication but also provide targets for improving N use efficiency in modern crops, balancing productivity with sustainability.

A conclusion of this study is that among biological processes commonly regulated by nitrate in Arabidopsis, Brachypodium and barley, molecular players of these processes can differ between species. These divergences highlight the importance of studying both model plants – to acquire a global view of involved processes and to serve as a basis for knowledge translation – and crops, to identify their specific molecular players.

More broadly, the pipeline developed here is not restricted to nitrate signaling or to the three species analyzed in this study. By first using Gene Ontology enrichment to identify potential conserved and species-specific biological processes, and then integrating orthology relationships to eliminate GO-annotation biases and examine regulation at the gene level, it provides a framework that can be transferred to any comparative transcriptomic study between species. A key advantage of this two-step strategy is that it reduces the risk of over-interpreting species-specific GO enrichment caused by uneven annotation quality: for example, “nitrate assimilation” GO appeared enriched in Arabidopsis and Brachypodium but not in barley in the GO per-species analysis, although the underlying orthologous genes were still responsive in barley, due to the lower GO annotation coverage in this species. Orthology analysis alone is insufficient for identifying candidate processes at large scale, while GO analysis alone can be biased by annotation differences between species. Our combined GO/OGp pipeline is therefore essential to correctly identify biological processes and candidate orthogroups of interest, although detailed analyses of specific gene families identified by this approach remain necessary to pursue functional validation.

## Supporting information

Spplemental Table 1

Spplemental Table 2

Spplemental Table 3A

Spplemental Table 3B

Spplemental Table 3C

Spplemental Table 4A

Spplemental Table 4B

Spplemental Table 4C

Spplemental Table 5

Spplemental Table 6

Spplemental Table 7

Spplemental Tables 8-12

## Acknowledgments

We thank Fabien Chardon for his help and ideas around the TopGO R package, Christian Meyer and the NUTS team for advices on experiments and analyses, Philippe Marechal and Hervé Ferry for help on plant culture and Sylvie Ferrario-Mery for proofreading. This work has benefited from the support of IJPB’s Plant Observatory platforms PO-Plants. This work was financed by the ANR project NiCe (ANR-17-CE20-0021). The POPS platform, IPS2 and IJPB benefit from the support of Saclay Plant Sciences-SPS (ANR-17-EUR-0007).

## Author contributions

MG, AK, MLM and TG conceived the experiments. MG and SP did the experiments. MG, VB and MLM did bioinformatic and statistical analyses. J.PT and LB generated OGps. MG and TG wrote the manuscript.

## Supplementary data

### Supplementary figures

**Supp Figure 1:** Schematic diagram of the experimental approach and RNAseq data extraction.

**Supp Figure 2:** Sentinel gene expression in response to 1mM NO_3_^-^ in roots of Arabidopsis (At), Brachypodium (Bd) and barley (Hv).

**Supp Figure 3:** Comparisons of DEG responses at 1.5h and 3h after nitrate treatment.

**Supp Figure 4:** Gene Ontology (GO) enrichment analysis for Arabidopsis, Brachypodium and barley after 1.5h of 1mM NO_3_^-^ treatment.

**Supp Figure 5:** Gene Ontology (GO) enrichment analysis for Arabidopsis, Brachypodium and barley after 3h of 1mM NO_3_^-^ treatment.

**Supp Figure 6**: Gene distributions in types of orthogroups.

**Supp Figure 7**: Brachypodium best 10 enriched biological process GO terms in response to 1mM nitrate treatment among Brachypodium/barley specific OGps.

**Supp Figure 8:** Gene Ontology (GO) comparative enrichment analysis for Arabidopsis, Brachypodium and barley after 1mM nitrate treatment (combination of 1.5h and 3h) for DEGs belonging to tri-species OGps.

### Supplementary tables

**Supp Table 1:** Sequences of primers

**Supp Table 2:** RNAseq data in response to 1.5h and 3h of nitrate treatment in Arabidopsis, Brachypodium and barley

**Supp Table 3A:** Significantly enriched biological process GO terms (combination of 1.5h and 3h DEG sets); per-species analyses

**Supp Table 3B:** Significantly enriched biological process GO terms (1.5h DEG sets); per-species analyses **Supp Table 3C:** Significantly enriched biological process GO terms (3h DEG sets); per-species analyses **Supp Table 4A:** Comparative GO enrichments analyses (combination of 1.5h and 3h DEG sets)

**Supp Table 4B:** Comparative GO enrichments analyses (1.5h DEG sets)

**Supp Table 4C:** Comparative GO enrichments analyses (3h DEG sets)

**Supp Table 5:** Composition of OGps in Arabidopsis, Brachypodium and barley genes

**Supp Table 6:** Significantly enriched biological process GO terms in DEGs considering Brachypodium/barley bi-species OGps (combination of 1.5h and 3h DEG sets); per-species analyses

**Supp Table 7:** Comparative GO enrichment analysis considering tri-species OGps (combination of 1.5h and 3h DEG sets)

**Supp Table 8:** Gene nitrate-response in tri-species OGps belonging to selected significantly enriched nitrate-related GO terms.

**Supp Table 9:** Gene nitrate-response in tri-species OGps belonging to selected significantly enriched nitrogen-related GO terms.

**Supp Table 10:** Gene nitrate-response in tri-species OGps belonging to “response to gibberellin” GO term.

**Supp Table 11:** Gene nitrate-response in tri-species OGps belonging to “rRNA processing” GO term.

**Supp Table 12:** Gene nitrate-response in tri-species OGps belonging to selected significantly enriched membrane trafficking and cell wall biogenesis GO terms.

## References

1. Alexa A, Rahnenführer J, Lengauer T. 2006. Improved scoring of functional groups from gene expression data by decorrelating GO graph structure. Bioinformatics 22, 1600–1607.

2. Alexa A, Rahnenfuhrer J. 2016. topGO: enrichment analysis for Gene Ontology. R Packag. Version 2.26.0. R Package version 2.26.0.

3. Alvarez M, Riveras E, Vidal EA, et al. 2014. Systems approach identifies TGA1 and TGA4 transcription factors as important regulatory components of the nitrate response of Arabidopsis thaliana roots. Plant Journal, 1–13.

4. Araus V, Vidal EA, Puelma T, Alamos S, Mieulet D, Guiderdoni E, Gutiérrez RA. 2016. Members of BTB gene family of scaffold proteins suppress nitrate uptake and nitrogen use efficiency. Plant Physiology 171, 1523–1532.

5. Berdy SE, Kudla J, Gruissem W, Gillaspy GE. 2001. Molecular Characterization of At5PTase1, an Inositol Phosphatase Capable of Terminating Inositol Trisphosphate Signaling. Plant Physiology 126, 801–810.

6. Bolger AM, Lohse M, Usadel B. 2014. Trimmomatic: A flexible trimmer for Illumina sequence data. Bioinformatics 30, 2114–2120.

7. Boycheva S, Dominguez A, Rolcik J, Boller T, Fitzpatrick TB. 2015. Consequences of a Deficit in Vitamin B6 Biosynthesis de Novo for Hormone Homeostasis and Root Development in Arabidopsis. Plant Physiology 167, 102–117.

8. Britto DT, Siddiqi MY, Glass ADM, Kronzucker HJ. 2001. Futile transmembrane NH 4 cycling : A cellular hypothesis to explain ammonium toxicity in plants. 98, 4255–4258.

9. Brooks MD, Cirrone J, Pasquino A V., et al. 2019. Network Walking charts transcriptional dynamics of nitrogen signaling by integrating validated and predicted genome-wide interactions. Nature Communications 10, 1569.

10. Camut L, Gallova B, Jilli L, et al. 2021. Nitrate signaling promotes plant growth by upregulating gibberellin biosynthesis and destabilization of DELLA proteins. Current Biology 31, 4971–4982.e4.

11. Canales J, Moyano TC, Villarroel E, Gutiérrez RA. 2014. Systems analysis of transcriptome data provides new hypotheses about Arabidopsis root response to nitrate treatments. Frontiers in Plant Science 5, 1–14.

12. Castaings L, Camargo A, Pocholle D, et al. 2009. The nodule inception-like protein 7 modulates nitrate sensing and metabolism in Arabidopsis. Plant Journal 57, 426–435.

13. Castle S, Randall P. 1987. Effects of Sulfur Deficiency on the Synthesis and Accumulation of Proteins in the Developing Wheat Seed. Australian Journal of Plant Physiology 14, 503.

14. Chardon F, Cue G, Delannoy E, Aub F. 2020. The Consequences of a Disruption in Cyto-Nuclear Coadaptation on the Molecular Response to a Nitrate Starvation in Arabidopsis. Plants.

15. Chen L, Ortiz-Lopez A, Jung A, Bush DR. 2001. ANT1, an aromatic and neutral amino acid transporter in Arabidopsis. Plant Physiology 125, 1813–1820.

16. Chu X, Wang JG, Li M, et al. 2021. HBI transcription factor-mediated ROS homeostasis regulates nitrate signal transduction. Plant Cell 33, 3004–3021.

17. Colinas M, Eisenhut M, Tohge T, Pesquera M, Fernie AR, Weber APM, Fitzpatrick TB. 2016. Balancing of B6 vitamers is essential for plant development and metabolism in arabidopsis. Plant Cell 28, 439–453.

18. David LC, Girin T, Fleurission E, Phommabouth E, Mahfoudhi A, Citerne S, Berquin P, Daniel-Vedele, Krapp A, Ferrario-Mery S. 2019. Developmental and physiological responses of Brachypodium distachyon to fluctuating nitrogen availability. Scientific Reports, 1–17.

19. Des Marais DL, Juenger TE. 2015. Brachypodium and the Abiotic Environment. In: Vogel JP, ed. Genetics and Genomics of Brachypodium. Springer International Publishing Switzerland, 291–311.

20. Dissanayake I, Rodriguez-medina J, Brady SM, Tanurdzic M. 2019. Transcriptional dynamics of bread wheat in response to nitrate and phosphate supply reveal functional divergence of genetic factors involved in nitrate and phosphate signaling. BioRxiv 551069; doi: 10.1101/551069.

21. Dobin A, Davis CA, Schlesinger F, Drenkow J, Zaleski C, Jha S, Batut P, Chaisson M, Gingeras TR. 2013. STAR: Ultrafast universal RNA-seq aligner. Bioinformatics 29, 15–21.

22. Du H, Ning L, He B, Wang Y, Ge M, Xu J, Zhao H. 2020. Cross-species root transcriptional network analysis highlights conserved modules in response to nitrate between maize and sorghum. International Journal of Molecular Sciences 21, 1–20.

23. Edgar R, Domrachev M, Lash AE. 2002. Gene Expression Omnibus: NCBI gene expression and hybridization array data repository. Nucleic Acids Research 30, 207–210.

24. Emms DM, Kelly S. 2019. OrthoFinder: Phylogenetic orthology inference for comparative genomics. Genome Biology 20.

25. Francia E, Morcia C, Pasquariello M, Mazzamurro V, Milc JA, Rizza F, Terzi V, Pecchioni N. 2016. Copy number variation at the HvCBF4–HvCBF2 genomic segment is a major component of frost resistance in barley. Plant Molecular Biology 92, 161–175.

26. Gagnot S, Tamby JP, Martin-Magniette ML, Bitton F, Taconnat L, Balzergue S, Aubourg S, Renou JP, Lecharny A, Brunaud V. 2008. CATdb: A public access to arabidopsis transcriptome data from the URGV-CATMA platform. Nucleic Acids Research 36, 986–990.

27. Gingerich DJ, Hanada K, Shiu SH, Vierstra RD. 2007. Large-scale, lineage-specific expansion of a bric-a-brac/tramtrack/broad complex ubiquitin-ligase gene family in rice. Plant Cell 19, 2329–2348.

28. Girin T, David LC, Chardin C, Sibout R, Krapp A, Ferrario-Méry S, Daniel-Vedele F. 2014. Brachypodium: A promising hub between model species and cereals. Journal of Experimental Botany 65, 5683–5686.

29. Golani Y, Kaye Y, Gilhar O, Ercetin M, Gillaspy G, Levine A. 2013. Inositol polyphosphate phosphatidylinositol 5-Phosphatase9 (At5PTase9) controls plant salt tolerance by regulating endocytosis. Molecular Plant 6, 1781–1794.

30. Guan P, Ripoll J-J, Wang R, Vuong L, Bailey-Steinitz LJ, Ye D, Crawford NM. 2017. Interacting TCP and NLP transcription factors control plant responses to nitrate availability. Proceedings of the National Academy of Sciences, 201615676.

31. Guan P, Wang R, Nacry P, Breton G, Kay SA, Pruneda-Paz JL, Davani A, Crawford NM. 2014. Nitrate foraging by *Arabidopsis* roots is mediated by the transcription factor TCP20 through the systemic signaling pathway. Proceedings of the National Academy of Sciences 111, 15267–15272.

32. Guo F, Huang Y, Qi P, Lian G, Hu X, Han N, Wang J, Zhu M, Qin Q, Bian H. 2020. Functional analysis of auxin receptor OsTIR1 / OsAFB family members in rice grain yield, tillering, plant height, root system, germination, and auxinic herbicide resistance.

33. Hanchi M, Thibaud M, Légeret B, et al. 2020. The Phosphate Fast-Responsive Genes PECP1 and PPsPase1 Affect Phosphocholine and Phosphoethanolamine Content. Plant Physiology, 176, 2943–2962.

34. Hardtke CS, Pacheco-Villalobos D. 2015. The Brachypodium distachyon Root System: A Tractable Model to Investigate Grass Roots. In: Vogel JP, ed. Genetics and Genomics of Brachypodium. Springer International Publishing Switzerland, 245–258.

35. Hesse H, Lipke J, Altmann T, Höfgen R. 1999. Molecular cloning and expression analyses of mitochondiral and plastidic isoforms of cystein synthase (O-acetylserine(thiol)lyase) from Arabidopsis thaliana. Amino Acids 16, 113–131.

36. Ho C, Lin S, Hu H, Tsay Y. 2009. CHL1 Functions as a Nitrate Sensor in Plants. Cell 138, 1184–1194.

37. Hochholdinger F, Park WJ, Sauer M, Woll K. 2004. From weeds to crops : genetic analysis of root development in cereals. Trends in Plant Science 9, 42–48.

38. Hu B, Jiang Z, Wang W, et al. 2019. Nitrate–NRT1.1B–SPX4 cascade integrates nitrogen and phosphorus signalling networks in plants. Nature Plants 5, 401–413.

39. Jia Q, Zhang J, Westcott S. 2009. GA-20 oxidase as a candidate for the semidwarf gene sdw1/denso in barley. Functional & Integrative Genomics 9, 255–262.

40. Kant S, Peng M, Rothstein SJ. 2011. Genetic Regulation by NLA and MicroRNA827 for Maintaining Nitrate-Dependent Phosphate Homeostasis in Arabidopsis. 7.

41. Kiba T, Feria-Bourrellier A-B, Lafouge F, et al. 2012. The Arabidopsis Nitrate Transporter NRT2 . 4 Plays a Double Role in Roots and Shoots of Nitrogen-Starved Plants. The Plant Cell 24, 245–258.

42. Kopylova E, Noé L, Touzet H. 2012. SortMeRNA: Fast and accurate filtering of ribosomal RNAs in metatranscriptomic data. Bioinformatics 28, 3211–3217.

43. Krapp A, Berthomé R, Orsel M, Boutet-Mercey S, Yu A, Castaings L, Elftieh S, Major H, Renou JP, Daniel-Vedele F. 2011. Arabidopsis Roots and Shoots Show Distinct Temporal Adaptation Patterns toward Nitrogen Starvation. Plant Physiology 157, 1255–1282.

44. Krouk G, Crawford NM, Coruzzi GM, Tsay YF. 2010*a*. Nitrate signaling: Adaptation to fluctuating environments. Current Opinion in Plant Biology 13, 266–273.

45. Krouk G, Mirowski P, LeCun Y, Shasha DE, Coruzzi GM. 2010*b*. Predictive network modeling of the high-resolution dynamic plant transcriptome in response to nitrate. Genome Biology 11, R123.

46. Lezhneva L, Kiba T, Feria-Bourrellier A, Lafouge F, Boutet-Mercey S, Zoufan P, Sakakibara H, Daniel-Vedele F, Krapp A. 2014. The Arabidopsis nitrate transporter NRT2.5 plays a role in nitrate acquisition and remobilization in nitrogen-starved plants. The Plant Journal 80, 230–241.

47. Liu K, Diener A, Lin Z, Liu C, Sheen J. 2020. Primary nitrate responses mediated by calcium signalling and diverse protein phosphorylation. Journal of Experimental Botany 71, 4428–4441.

48. Liu K, Niu Y, Konishi M, et al. 2017. Discovery of nitrate–CPK–NLP signalling in central nutrient–growth networks. Nature, 1–23.

49. Liu K-H, Liu M, Lin Z, et al. 2022. NIN-like protein 7 transcription factor is a plant nitrate sensor. Science 377, 1419–1425.

50. Loqué D, von Wirén N. 2004. Regulatory levels for the transport of ammonium in plant roots. Journal of Experimental Botany 55, 1293–1305.

51. Loqué D, Yuan L, Kojima S, Gojon A, Wirth J, Gazzarrini S, Ishiyama K, Takahashi H, von Wirén N. 2006. Additive contribution of AMT1 ; 1 and AMT1 ; 3 to high-affinity ammonium uptake across the plasma membrane of nitrogen-deficient Arabidopsis roots. the plant journal, 522–534.

52. Lunn JE, Delorge I, Figueroa CM, Van Dijck P, Stitt M. 2014. Trehalose metabolism in plants. Plant Journal 79, 544–567.

53. Lycett G. 2008. The role of Rab GTPases in cell wall metabolism. Journal of Experimental Botany 59, 4061–4074.

54. Maeda Y, Konishi M, Kiba T, Sakuraba Y, Sawaki N, Kurai T, Ueda Y, Sakakibara H, Yanagisawa S. 2018. A NIGT-centred transcriptional cascade regulates Nitrate signalling and incorporates phosphorus starvation signals in Arabidopsis. Nature Communications.

55. Mao J, Tian Z, Sun J, Wang D, Yu Y, Li S. 2025. The crosstalk between nitrate signaling and other signaling molecules in Arabidopsis thaliana. Frontiers in Plant Science 16.

56. Marchive C, Roudier F, Castaings L, Bréhaut V, Blondet E, Colot V, Meyer C, Krapp A. 2013. Nuclear retention of the transcription factor NLP7 orchestrates the early response to nitrate in plants. Nature Communications 4, 1713.

57. Marschner H, Römheld V. 1994. Strategies of plants for acquisition of iron. Plant and Soil, 261–274.

58. McCarthy DJ, Chen Y, Smyth GK. 2012. Differential expression analysis of multifactor RNA-Seq experiments with respect to biological variation. Nucleic Acids Research 40, 4288–4297.

59. Medici A, Marshall-colon A, Ronzier E, Szponarski W, Wang R, Gojon A, Crawford NM, Ruffel S, Coruzzi GM, Krouk G. 2015. AtNIGT1/HRS1 integrates nitrate and phosphate signals at the Arabidopsis root tip. Nature Communications 6, 1–11.

60. Medici A, Szponarski W, Dangeville P, Safi A. 2019. Identification of Molecular Integrators Show that Nitrogen Actively Controls Response in Plants the Phosphate Starvation. Plant Cell.

61. Miyawaki K, Matsumoto-kitano M, Kakimoto T. 2004. Expression of cytokinin biosynthetic isopentenyltransferase genes in Arabidopsis : tissue speci ® city and regulation by auxin, cytokinin, and nitrate., 128–138.

62. Naulin PA, Armijo GI, Vega AS, Tamayo KP, Gras DE, Cruz J De, Guti RA. 2020. Nitrate Induction of Primary Root Growth Requires Cytokinin Signaling in Arabidopsis thaliana. Plant Cell Physiology, 61, 342–352.

63. Obertello M, Shrivastava S, Katari MS, Coruzzi GM. 2015. Cross-species network analysis uncovers conserved nitrogen-regulated network modules in rice. Plant Physiology 168, 1830–1843.

64. Osmont KS, Sibout R, Hardtke CS. 2007. Hidden Branches : Developments in Root System Architecture, Annual review of plant biology, 58, 93–113.

65. Pacheco-villalobos D, Sankar M, Ljung K, Hardtke CS. 2013. Disturbed Local Auxin Homeostasis Enhances Cellular Anisotropy and Reveals Alternative Wiring of Auxin- ethylene Crosstalk in Brachypodium distachyon Seminal Roots. PLOS 9.

66. Plett D, Toubia J, Garnett T, Tester M, Kaiser BN, Baumann U. 2010. Dichotomy in the NRT gene families of dicots and grass species. PLoS ONE 5.

67. Rieu I, Powers SJ. 2009. Real-Time Quantitative RT-PCR: Design, Calculations, and Statistics. The Plant Cell 21, 1031–1033.

68. Rieu I, Ruiz-Rivero O, Fernandez-Garcia N, et al. 2008. The gibberellin biosynthetic genes AtGA20ox1 and AtGA20ox2 act, partially redundantly, to promote growth and development throughout the Arabidopsis life cycle. Plant Journal 53, 488–504.

69. Rigaill G, Balzergue S, Brunaud V, et al. 2018. Synthetic data sets for the identification of key ingredients for RNA-seq differential analysis. Briefings in Bioinformatics 19, 65–76.

70. Riveras E, Alvarez JM, Vidal E a., Oses C, Vega A, Gutiérrez RA. 2015. The Calcium Ion Is a Second Messenger in the Nitrate Signaling Pathway of Arabidopsis. Plant Physiology 169, 1397–1404.

71. Robert HS, Quint A, Brand D, Vivian-Smith A, Offringa R. 2009. BTB and TAZ domain scaffold proteins perform a crucial function in Arabidopsis development. Plant Journal 58, 109–121.

72. Rubin G, Tohge T, Matsuda F, Saito K, Sheible W. 2009. Members of the LBD Family of Transcription Factors Repress Anthocyanin Synthesis and Affect Additional Nitrogen Responses in Arabidopsis. The Plant Cell 21, 3567–3584.

73. Ruffel S, Del Rosario J, Lacombe B, Rouached H, Gutiérrez RA, Coruzzi GM, Krouk G. 2025. Nitrate Sensing and Signaling in Plants: Comparative Insights and Nutritional Interactions. Annual Review of Plant Biology 76, 25–52.

74. Scheible WR, Morcuende R, Czechowski T, Fritz C, Osuna D, Palacios-Rojas N, Schindelasch D, Thimm O, Udvardi MK, Stitt M. 2004. Genome-wide reprogramming of primary and secondary metabolism, protein synthesis, cellular growth processes, and the regulatory infrastructure of arabidopsis in response to nitrogen. Plant Physiology 136, 2483–2499.

75. Shahzad Z, Amtmann A. 2017. Food for thought: how nutrients regulate root system architecture. Current Opinion in Plant Biology 39, 80–87.

76. Smith S, de Smet I. 2012. Root system architecture: Insights from Arabidopsis and cereal crops. Philosophical Transactions of the Royal Society B: Biological Sciences 367, 1441–1452.

77. Tian Q, Sun P, Zhang W. 2009. Ethylene is involved in nitrate-dependent root growth and branching in Arabidopsis thaliana. New Phytologist 184, 918–931.

78. Tripathy MK, Deswal R, Sopory SK. 2021. Plant RABs: Role in Development and in Abiotic and Biotic Stress Responses. Current Genomics 22, 26–40.

79. Varala K, Marshall-colón A, Cirrone J, Brooks MD, Pasquino AV, Léran S, Mittal S, Rock TM, Edwards MB, Kim GJ, Ruffel S, McCombie WR, Shasha D, Coruzzi GM. 2018. Temporal transcriptional logic of dynamic regulatory networks underlying nitrogen signaling and use in plants. Proceedings of the National Academy of Sciences of the United States of America 115, 6494–6499.

80. Vidal E A, Araus V, Lu C, Parry G, Green PJ, Coruzzi GM, Gutiérrez R. 2010. Nitrate-responsive miR393/AFB3 regulatory module controls root system architecture in Arabidopsis thaliana. Proceedings of the National Academy of Sciences of the United States of America 107, 4477–82.

81. Vidal EA, Alvarez JM, Araus V, Riveras E, Brooks MD, Krouk G, Ruffel S, Lejay L, Crawford NM, Coruzzi GM, Gutiérrez RA. 2020. Nitrate in 2020: Thirty years from transport to signaling networks. Plant Cell 32, 2094–2119.

82. Woollard AAD, Moore I. 2008. The functions of Rab GTPases in plant membrane traffic. Current Opinion in Plant Biology, 610–619.

83. Xu G, Fan X, Miller AJ. 2012. Plant Nitrogen Assimilation and Use Efficiency. Annual reviews, 63:153.

84. Xu N, Wang R, Zhao L, et al. 2016. The Arabidopsis NRG2 Protein Mediates Nitrate Signaling and Interacts with and Regulates Key Nitrate Regulators. The Plant Cell 28, 485–504.

