## Supplemental Figures for "Comparative cross-species transcriptomic analysis identifies new candidates of Pooideae nitrate response"

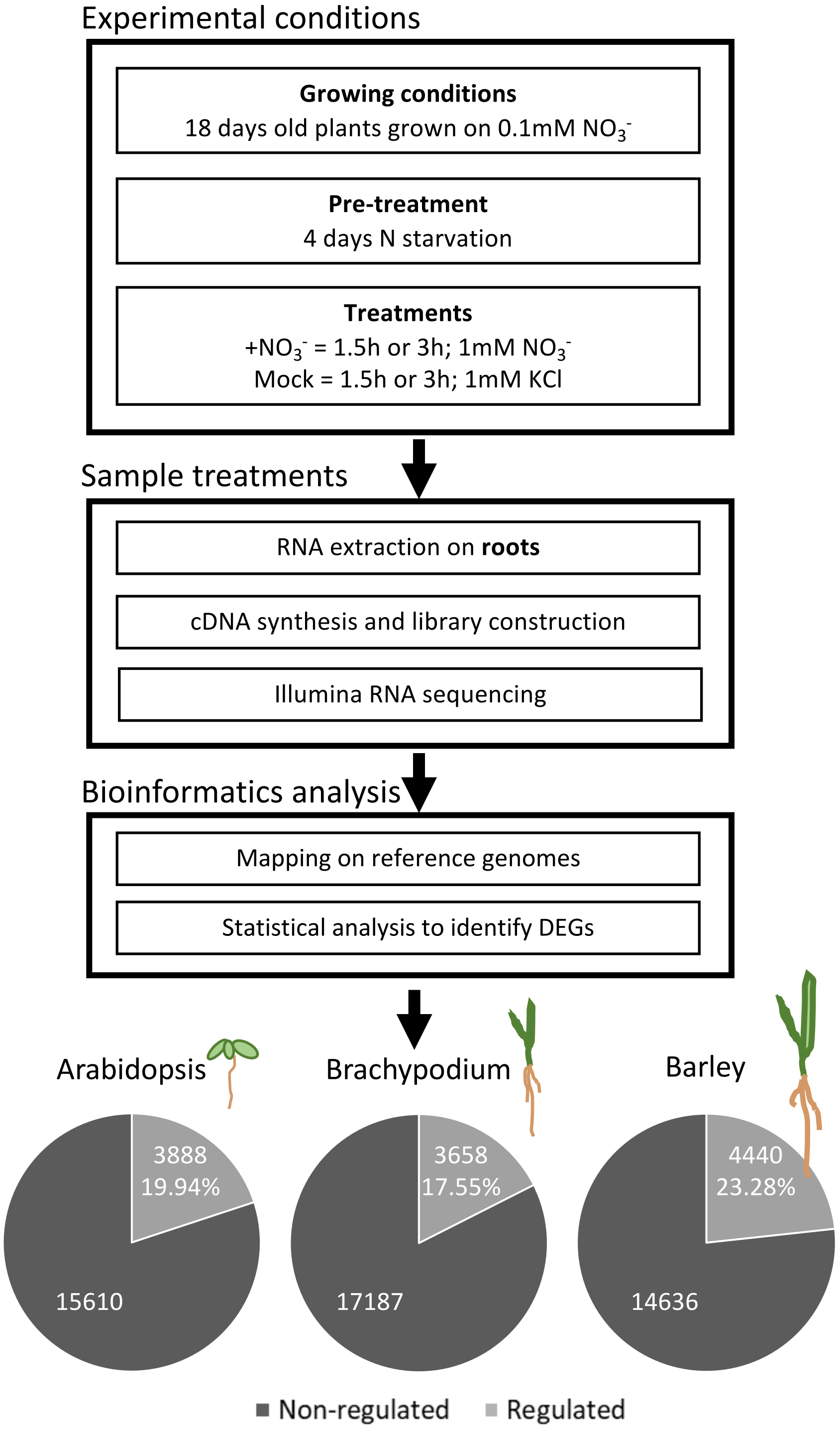


**Supplementary Figure 1: Schematic diagram of the experimental approach and RNAseq data extraction.** Arabidopsis, Brachypodium and barley plantlets were grown hydroponically for 18 days on basal medium. After 4 days of nitrogen starvation, plantlets were treated for 1.5h or 3h with 1mM $NO_3^-$ or 1mM KCl (mock-treatment). Roots were collected and RNA were extracted and processed for RNAseq. Mapping and differential analysis finally led to the identification of Differentially Expressed Genes (DEGs) between nitrate- and mock-treated samples for each timepoint and each species. The count of nitrate-regulated genes (combination of 1.5h and 3h sets) and their proportion among detected genes are represented by pie charts.

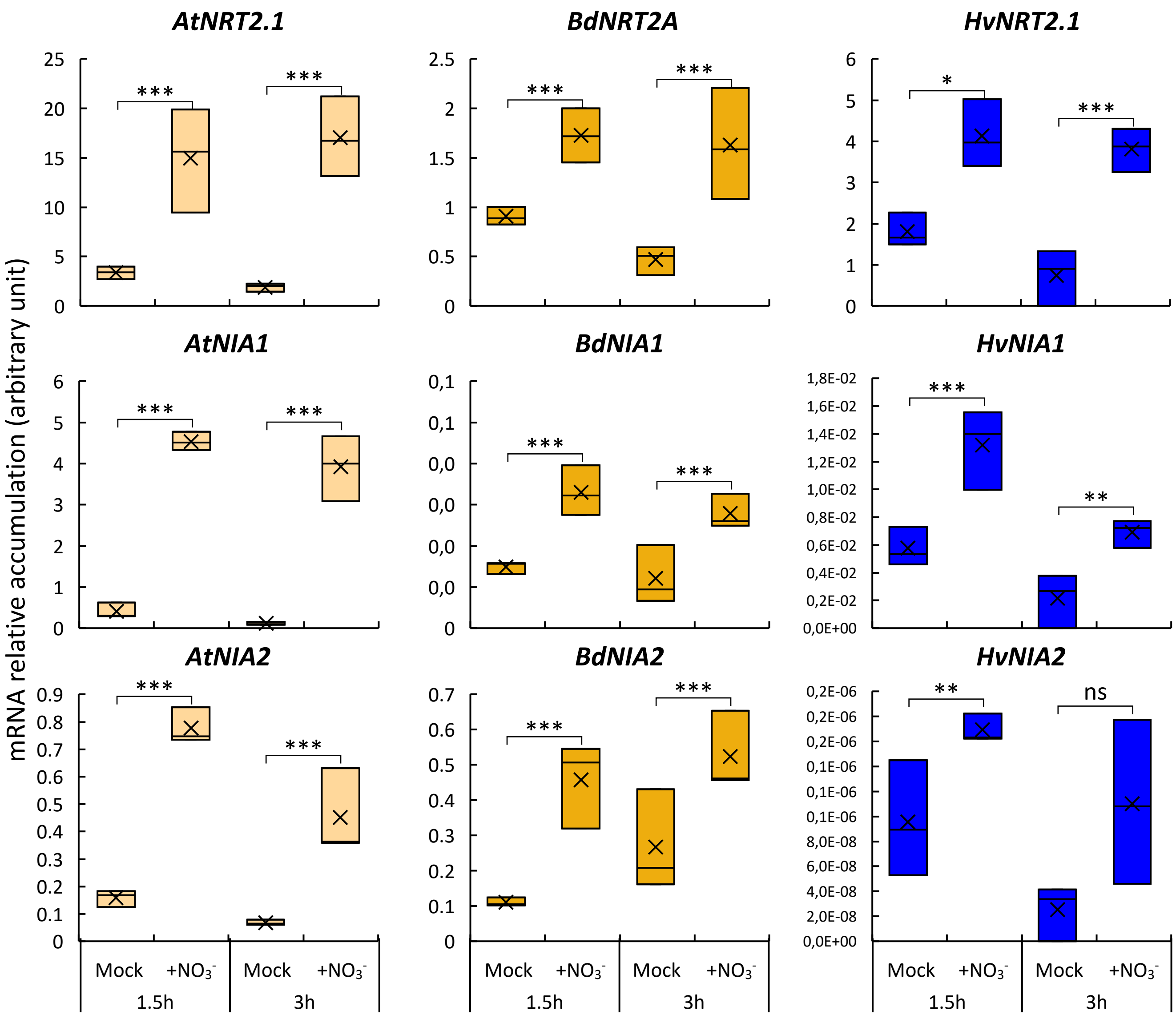


**Supplementary Figure 2: Sentinel gene expression in response to 1mM $NO_3^-$ in roots of Arabidopsis (At), Brachypodium (Bd) and barley (Hv).** RT-qPCR have been performed on the samples from the transcriptomics experiment. The boxplots represent minimum, median, mean (cross) and maximum values. Stars (*) represent the statistical significance between nitrate-treated samples (+$NO_3^-$) and their corresponding mock sample by a non-parametric ANOVA test (*pvalue<0.05; **pvalue<0.01; ***pvalue<0.001; ns non-significant; n=3).

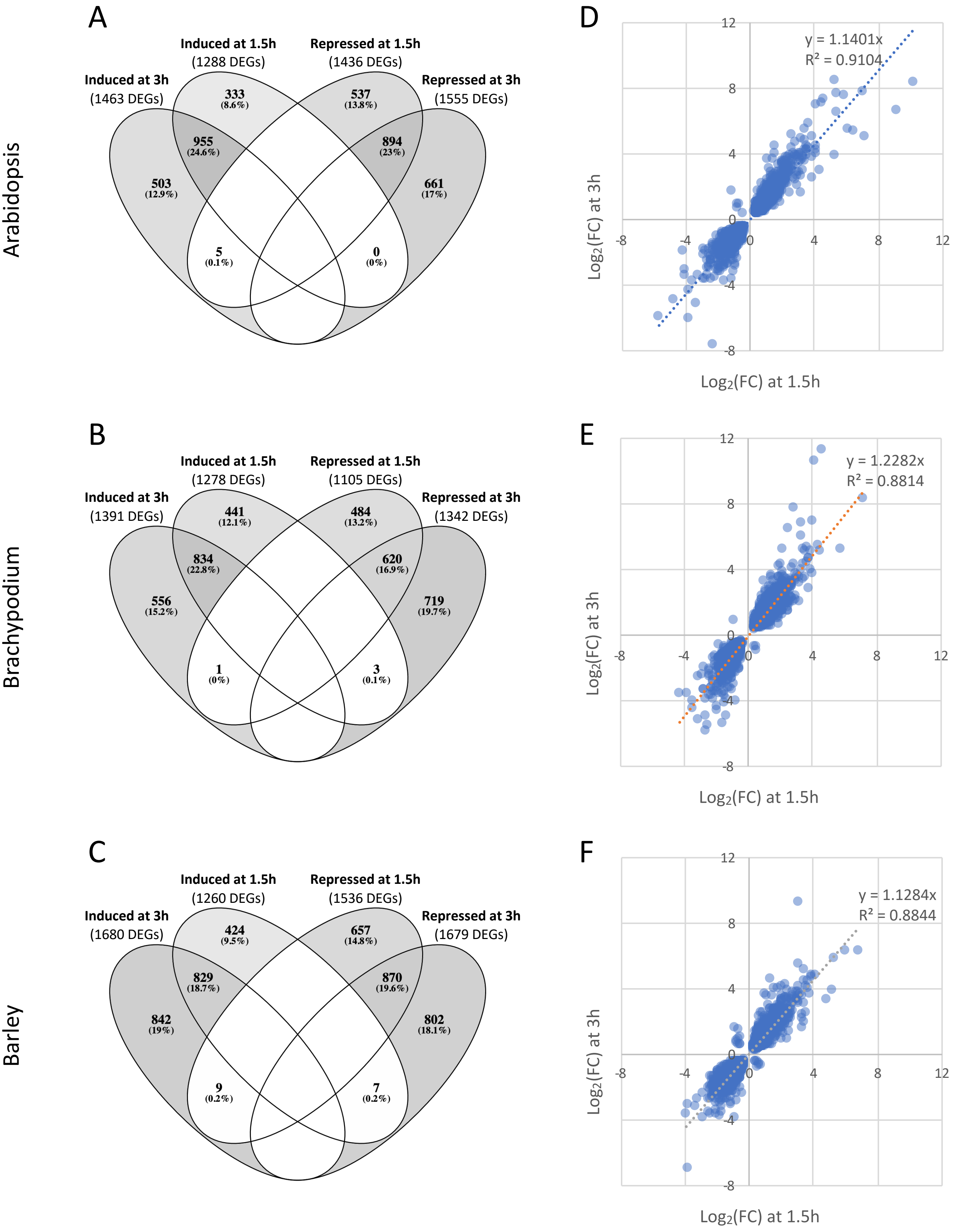


**Supplementary Figure 3: Comparisons of DEG responses at 1.5h and 3h after nitrate treatment. (A-C)** Venn diagrams of induced and repressed DEGs at 1.5h and 3h after addition of nitrate in Arabidopsis **(A)**, Brachypodium **(B)** and barley **(C)**. **(D-F)** Correlations of $Log_2$(FC) at 1.5h and 3h for DEGs significantly responding at both timepoints in Arabidopsis (**D**; 1854 genes), Brachypodium (**E**; 1458 genes) and barley (**C**; 1715 genes). FC = expression Fold Change between nitrate-treated and mock conditions. Equation and $R^2$ of linear regression are shown. Note that the slopes of the linear regressions are higher than 1, indicating a global higher response at  3h post-treatment.

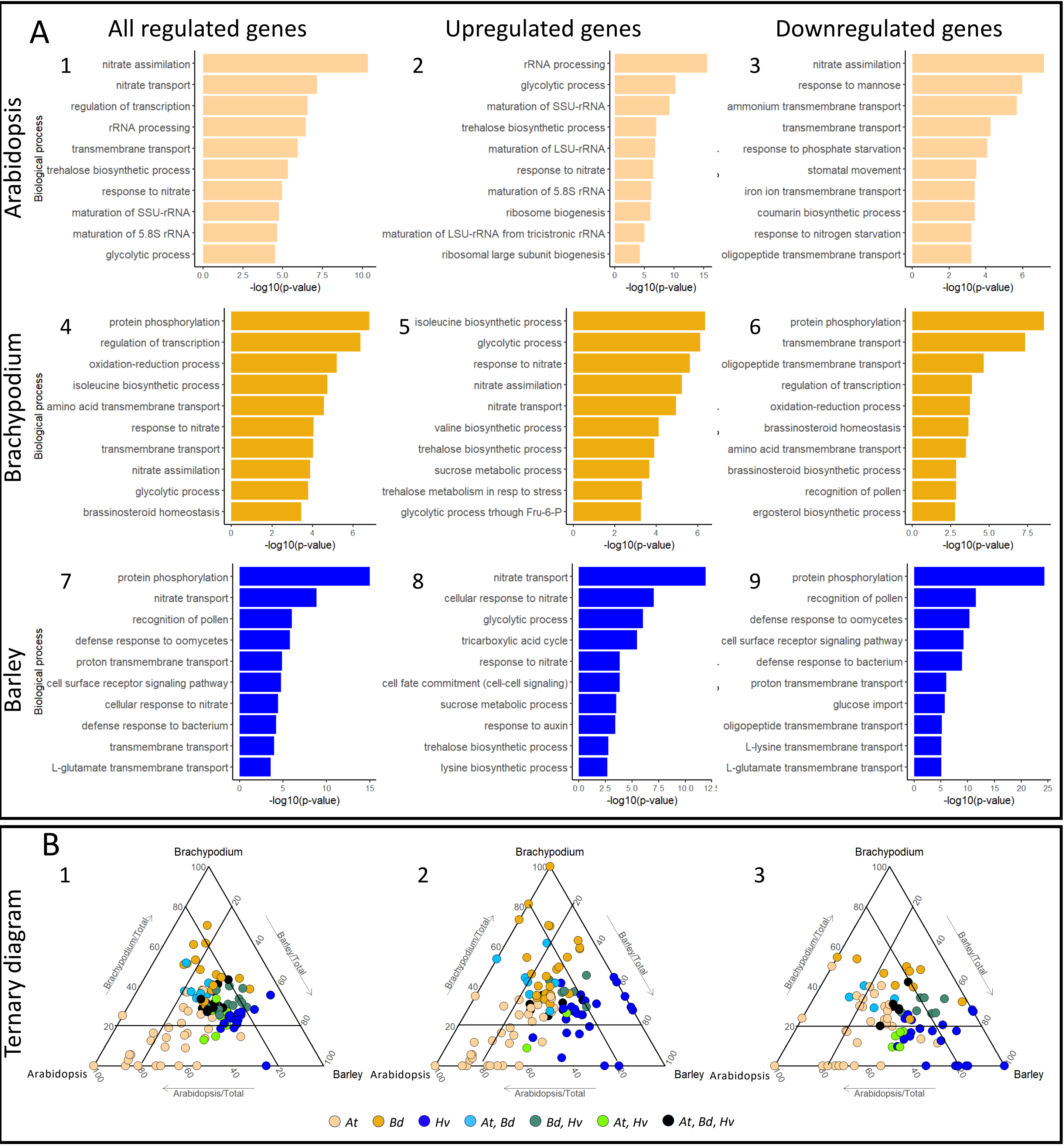


**Supplementary Figure 4: Gene Ontology (GO) enrichment analysis for Arabidopsis, Brachypodium and barley after 1.5h of 1mM $NO_3^-$ treatment. (A)** The 10 best enriched GO terms (based on their p-value score) were plotted for each species. The full lists of enriched GO terms are available in Supp Table 3B. **(B)** Ternary diagrams representing the distribution of GOs that belong to the 50 most-significantly enriched GO terms in at least one species (112, 108 and 89 GO terms for B1, B2 and B3, respectively). Each axes represent the Normalized Enrichment Score (NES) for one species and each dot represents a GO term. Colors indicate for which species the GO term was significantly enriched in initial per-species analysis (ElimFisher test, pval<0.05). Internal bars represent a 20% threshold which is used for comparing enrichment scores between species (see Results). The lists of selected GOs and their NES are available in Supplementary Table 4B. For both (A) and (B), the first column represents biological process GO terms considering all the DEGs; the second column represents biological process GO terms considering only the upregulated DEGs; the third column, considering only downregulated DEGs. Gene ontology terms, their enrichment scores and their p-values were generated using topGO R package.

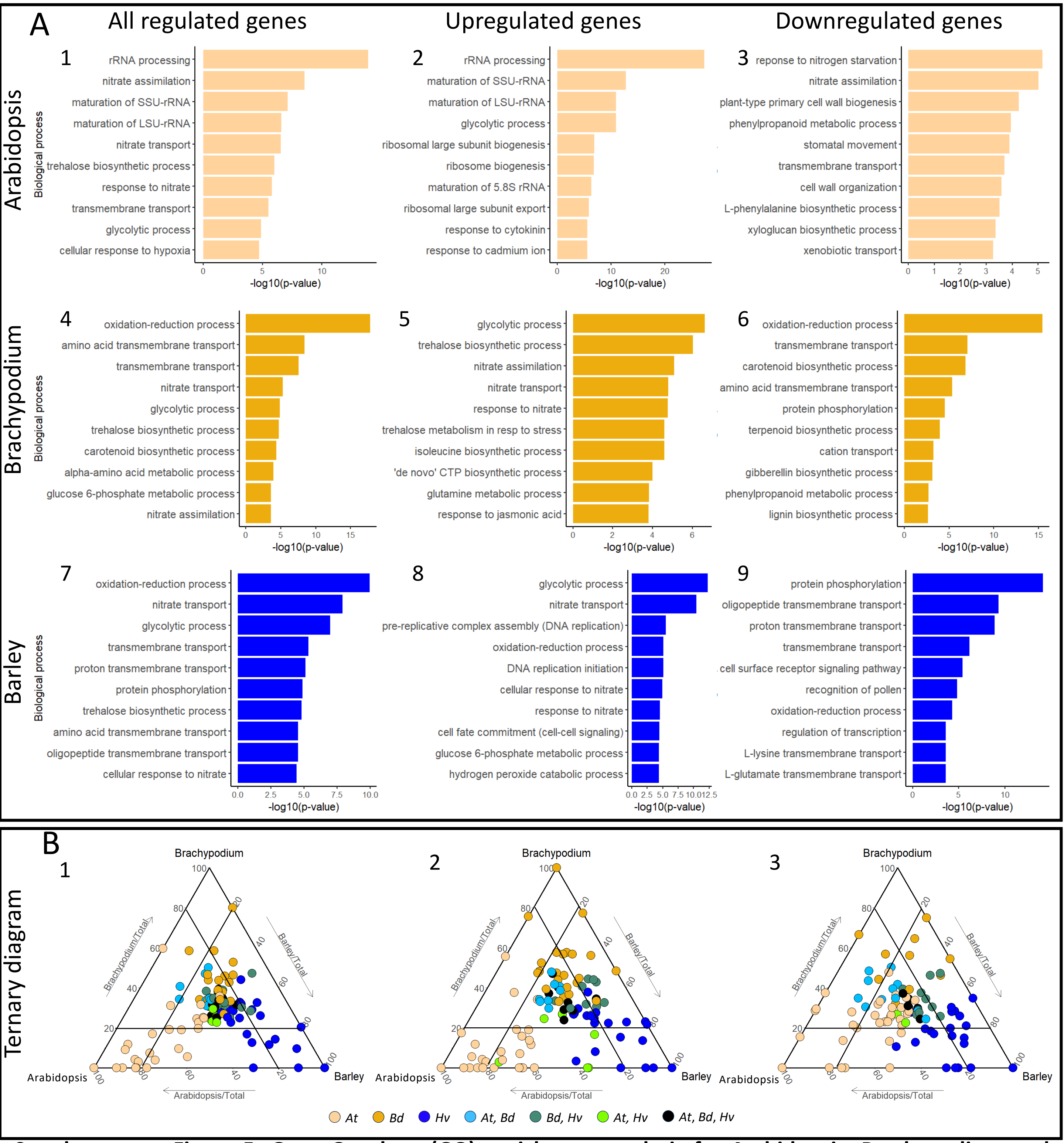


**Supplementary Figure 5: Gene Ontology (GO) enrichment analysis for Arabidopsis, Brachypodium and barley after 3h of 1mM $NO_3^-$ treatment. (A)** The 10 best enriched GO terms (based on their p-value score) were plotted for each species. The full lists of enriched GO terms are available in Supp Table 3C. **(B)** Ternary diagrams representing the distribution of GOs that belong to the 50 most-significantly enriched GO terms in at least one species (114, 115 and 93 GO terms for B1, B2 and B3, respectively). Each axes represent the Normalized Enrichment Score (NES) for one species and each dot represents a GO term. Colors indicate for which species the GO term was significantly enriched in initial per-species analysis (ElimFisher test, pval<0.05). Internal bars represent a 20% threshold which is used for comparing enrichment scores between species (see Results). The lists of selected GOs and their NES are available in Supp Table 4C. For both (A) and (B), the first column represents biological process GO terms considering all the DEGs; the second column represents biological process GO terms considering only the upregulated DEGs; the third column, considering only downregulated DEGs. Gene ontology terms, their enrichment scores and their p-values were generated using topGO R package.

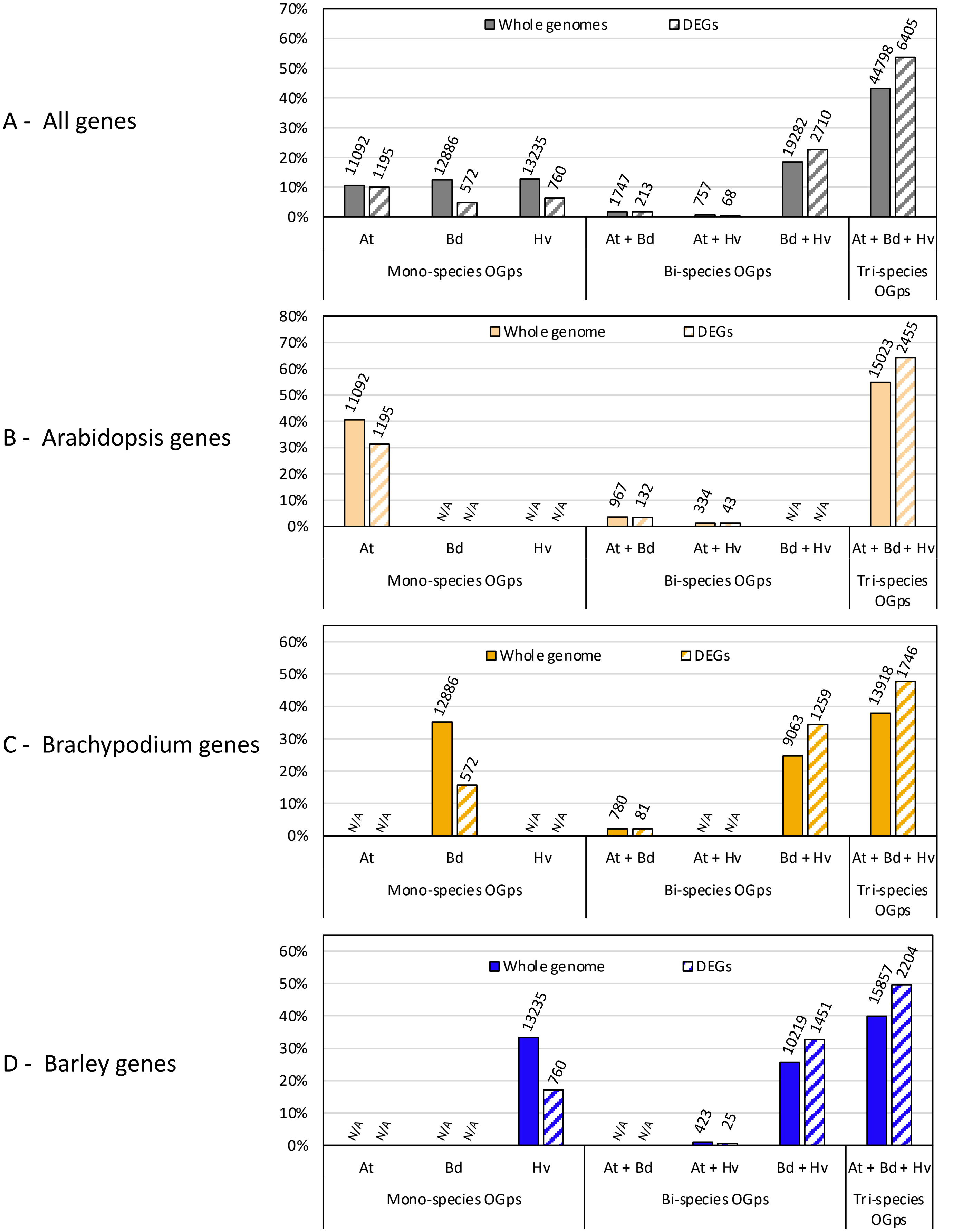


**Supplementary Figure 6**: **Gene distributions in types of orthogroups**. Distributions in OGps types for whole-genome gene models (plain bars) and DEGs (hatched bars) are presented. Percentages are relative to the total number of gene models or DEGs (A, 103797 and 11923, respectively; B, 27416 and 3825; C, 36647 and 3658; D, 39734 and 4440). Numbers of genes are indicated above each category. At (Arabidopsis), Bd (Brachypodium) and Hv (barley) indicate the species encompassed by the OGp types. For each graph, the distribution of DEGs is significantly different from the distribution of whole-genome gene models (chi-squared, pval<0.001).

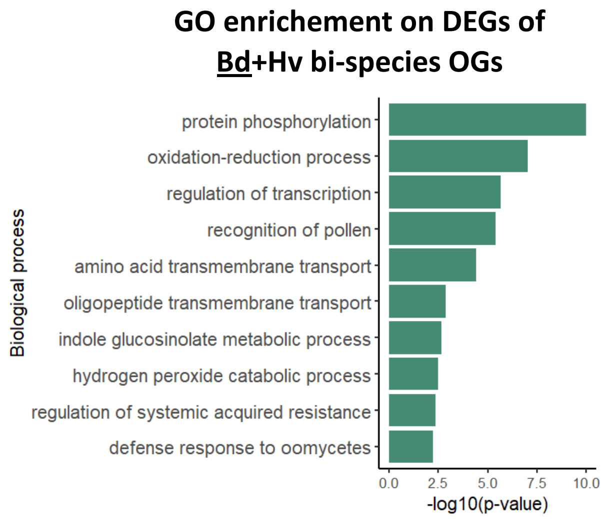


**Supplementary Figure 7: Brachypodium best 10 enriched biological process GO terms in response to 1mM nitrate treatment among Brachypodium-barley specific OGps.** Using topGO R package, significant enriched GO terms were identify on Brachypodium DEG lists belonging to Bd+Hv bi-species OGps. All significant enriched GO terms are available in Supp Table 6A. Analysis was also done on barley DEG lists belonging to Bd+Hv bi-species OGps, giving similar results (Supp Table 6B).

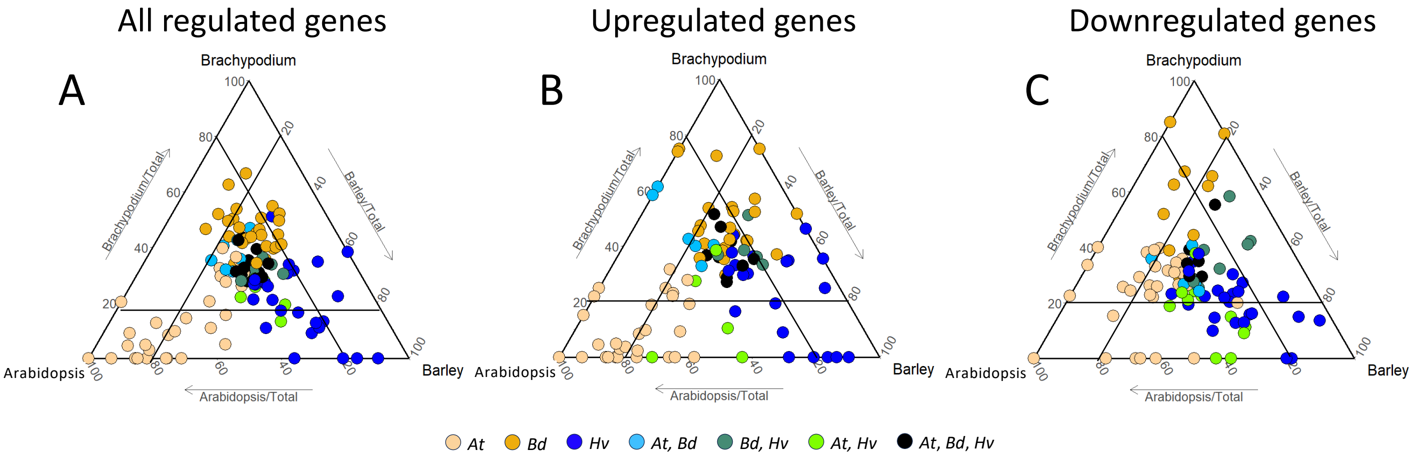


**Supplementary Figure 8: Gene Ontology (GO) comparative enrichment analysis for Arabidopsis, Brachypodium and barley after 1mM nitrate treatment (combination of 1.5h and 3h) for DEGs belonging to tri-species OGps.** Ternary diagrams represent the distribution of the 50 best enriched GO terms per species (giving totals of 109, 93, 84 GO terms for A, B and C, respectively, due to overlap between the species). Each axes represent the Normalized Enrichment Score for one species and each point represents a GO term. Colors indicate for which species the GO term was significantly enriched in initial per-species analysis (ElimFisher test, pval<0,05). Internal bars represent a 20% threshold which is used for comparing enrichment scores between species. **(A)** biological process GO terms considering all the DEGs from tri-species OGps; **(B)** biological process GO terms considering only the upregulated DEGs; **(C)** biological process GO terms considering only downregulated DEGs. Gene ontology terms, their enrichment scores and their p-values were generated using topGO R package. Complete data lists are available in Supp Table 7.
