## Supplementary material for "Comparative cross-species transcriptomic analysis identifies new candidates of Pooideae nitrate response": Spplemental Tables 8-12

A

| OGp | Arabidopsis gene names | *At* genes | | | | *Bd* genes | | | | | *Hv* genes | | | |
|---|---|---|---|---|---|---|---|---|---|---|---|---|---|---|
| OG0001808 | NRT1/ PTR FAMILY (NRT1.1/NPF6.3) | 2.3 | | | | | 4.06 | 4.29 | | | 5.16 | 2.2 | | |
| OG0002924 | BTB AND TAZ DOMAIN PROTEIN (BT1) (BT2) | 4.8 | 4.42 | | | 4 | | | | | 3.83 | 3.57 | 3.3 | |
| OG0004755 | NITRATE TRANSPORTER 3.1 (NRT3.1) | 2.06 | | | | 1.18 | 0.89 | | | | 2.31 | 1.11 | | |
| OG0007111 | CHLORIDE CHANNEL (CLCA) (CLCB) | 1.07 | 1.17 | | | 1.85 | | | | | 3.31 | | | |
| OG0010222 | NITRITE REDUCTASE 1 (NIR1) | 3.07 | | | | 2.89 | | | | | 3.39 | | | |
| OG0008828 | NITRATE TRANSPORTER2.5 (NRT2.5) | -3.63 | | | | -1.69 | | | | | -2.76 | | | |
| OG0000527 | NITRATE TRANSPORTER 2 (NRT2.1) (NRT2.2) (NRT2.4) | 2.49 | 2.3 | -1.57 | | | 1.58 | 2.23 | | | 2.99 | 2.61 | 2.75 | 1.11 |
| OG0002517 | NIN-LIKE PROTEIN (NLP6) (NLP7) | | -0.61 | | | 0.6 | | | | | -0.49 | | | |
| OG0009715 | ECTOPIC ROOT HAIR (ERH1) | -0.51 | | | | -0.63 | | | | | | | | |
| OG0002316 | DUF630 family protein, putative (DUF630 and DUF632) | | | | | | 0.85 | | | | | 1.2 | | |
| OG0005474 | AUXIN SIGNALING F-BOX (AFB3) (AFB2) | 0.57 | | | | | | | | | | | | |
| OG0007055 | NRT1/ PTR FAMILY 6.4 (NPF6.4) | -2.81 | | | | | | | | | | | | |
| OG0005537 | NRT1/ PTR FAMILY (NPF7.3/NRT1.5) (NPF7.2/NRT1.8) | 0.7 | -1.21 | | | | | | | | | | | |
| OG0005753 | NITRATE REGULATORY GENE 2 (NRG2) | | | | | 0.58 | | | | | | | | |
| OG0010661 | benzoyl-CoA reductase subunit C, putative (DUF630 and DUF632) | | | | | 0.35 | | | | | | | | |
| OG0009051 | NITROGEN LIMITATION ADAPTATION (NLA) | | | | | | | | | | 0.77 | | | |
| OG0004093 | DNA-directed RNA polymerase subunit beta, putative (DUF630 and DUF632) | | | | | | | | | | -0.39 | 0.58 | | |
| OG0003933 | kinesin-like protein ; DNA ligase (DUF630 and DUF632) | | | | | | | | | | | | | |
| OG0004654 | bZIP transcription factor, putative (DUF630 and DUF632) | | | | | | | | | | | | | |
| OG0006399 | DUF630 family protein (DUF630 and DUF632) | | | | | | | | | | | | | |
| OG0010033 | bZIP transcription factor (DUF630 and DUF632) | | | | | | | | | | | | | |
| OG0011275 | ALTERED PHOSPHATE STARVATION RESPONSE 1 (APSR1) | | | | | | | | | | | | | |
| | Up | 32% | | | | 39% | | | | | 46% | | | |
| | Down | 18% | | | | 6% | | | | | 9% | | | |
| | Total | 50% | | | | 45% | | | | | 54% | | | |

B

| OGp | Arabidopsis gene names | *At* genes | | | | *Bd* genes | | | | | *Hv* genes | | | |
|---|---|---|---|---|---|---|---|---|---|---|---|---|---|---|
| OG0001808 | NRT1/PTR FAMILY 6.3 (NRT1.1/NPF6.3) | 2.3 | | | | | 4.06 | 4.29 | | | 5.16 | 2.2 | | |
| OG0004755 | NITRATE TRANSPORTER 3.1 (NRT3.1) | 2.06 | | | | 1.18 | 0.89 | | | | 2.31 | 1.11 | | |
| OG0007111 | CHLORIDE CHANNEL (CLCA) (CLCB) | 1.07 | 1.17 | | | 1.85 | | | | | 3.31 | | | |
| OG0003827 | NRT1/ PTR FAMILY (NPF1.2/NRT1.11) (NRT1.12) | | | -0.99 | | -1.75 | | | | | -2 | | | |
| OG0008828 | NITRATE TRANSPORTER 2.5 (NRT2.5) | -3.63 | | | | -1.69 | | | | | -2.76 | | | |
| OG0000527 | NITRATE TRANSPORTER 2 (NRT2.1) (NRT2.2) (NRT2.4) | 2.49 | 2.3 | -1.57 | | | 1.58 | 2.23 | | | 2.99 | 2.61 | 2.75 | 1.11 |
| OG0010011 | PEPTIDE TRANSPORTER 2 (PTR2) | | | | | -0.36 | | | | | -0.66 | | | |
| OG0012734 | C-TERMINALLY ENCODED PEPTIDE RECEPTOR 1 (CEPR1) | | | | | -1.99 | | | | | -1.46 | | | |
| OG0003595 | NRT1/ PTR FAMILY 2.12 (NPF2.12/NRT1.6) | | -0.49 | | | | | | | | | | | |
| OG0007055 | NRT1/ PTR FAMILY 6.4 (NPF6.4) | -2.81 | | | | | | | | | | | | |
| OG0005537 | NRT1/ PTR FAMILY 7.3 (ATNPF7.3/NRT1.5) | 0.7 | -1.21 | | | | | | | | | | | |
| OG0005753 | NITRATE REGULATORY GENE 2 (NRG2) | | | | | 0.58 | | | | | | | | |
| OG0009374 | C-TERMINALLY ENCODED PEPTIDE RECEPTOR 2 (CEPR2) | | | | | -1.63 | | | | | | | | |
| OG0001006 | COPPER AMINE OXIDASE ALPHA | | | | | | | | | | | | | |
| | Up | 28% | | | | 36% | | | | | 38% | | | |
| | Down | 24% | | | | 23% | | | | | 17% | | | |
| | Total | 52% | | | | 59% | | | | | 54% | | | |

**Supplementary Table 8: Gene nitrate-response in tri-species OGps belonging to selected significantly enriched nitrate-related GO terms.** GO terms “response to nitrate” [GO:0010167] **(A)** and “nitrate transport” [GO:0015706] **(B)** were identified as common to Arabidopsis, Brachypodium and barley by the comparative analysis on tri-species OGps. These GO fully encompass the enriched GO term “cellular response to nitrate” [GO:0071249] (OGps with [*]). Genes associated to those GO terms and belonging to tri-species OGps were extracted. Lines represent OGp; colored rectangles represent genes, grouped by species (*At*, Arabidopsis; *Bd*, Brachypodium; *Hv*, barley). Color scale and numbers indicate maximum $Log_2FC$ values (1.5h or 3h nitrate *vs* mock); blue, upregulated; red, downregulated; grey, not-regulated/not-detected. Proportions of upregulated, downregulated and total DEGs among represented genes are indicated below the tables for each species.

A

| OGp | Arabidopsis gene names | *At* genes | | | | *Bd* genes | | *Hv* genes |
|---|---|---|---|---|---|---|---|---|
| OGp0012489 | *DEGRADATION OF UREA 3 (DUR3)* | -2.25 | | | | -2.88 | | -2.69 |
| OGp0012092 | *ALLANTOINASE (ALN)* | -1.22 | | | | -1.56 | | -0.65 |
| OGp0011214 | *AUTOPHAGY 8H (APG8H/ATG8I)* | -0.50 | | | | -0.53 | | -0.50 |
| OGp0001713 | *CATALASE (CAT3/SEN2) (CAT1) (CAT2)* | -0.60 | | 1.12 | | 0.68 | 2.87 | 1.11 |
| OGp0006194 | *AUTOPHAGY 8C (ATG8C)* | -0.58 | | | | | | -0.49 |
| OGp0007681 | *GENERAL REGULATORY FACTOR 3 (GRF3)* | | | | | | 0.66 | 0.54 |
| OGp0004315 | *AUTOPHAGY 10 (ATG10)* | | | | | -0.56 | | |
| OGp0006772 | *SUGAR WILL EVENTUALLY BE EXPORTED (SWEET16) (SWEET17)* | | | | | | | -0.77 |
| OGp0007236 | *ABC1-LIKE KINASE RELATED TO CHLOROPHYLL DEGRADATION AND OXIDATIVE STRESS 1 (ACDO1)* | | | | | | | -0.45 |
| OGp0011120 | *AUTOPHAGY 6 (ATG6)* | | | | | | | -0.40 |
| OGp0012333 | *AUTOPHAGY 5 (ATG5)* | | | | | | | |
| OGp0012948 | *AUTOPHAGY 7 (ATG7)* | | | | | | | |
| OGp0001006 | *COPPER AMINE OXIDASE ALPHA (CUAO α ) (CUAOα2) (CUAOα3) (CUAOα)* | | | | | | | |
| | Up | 5% | | | | 19% | | 10% |
| | Down | 26% | | | | 25% | | 35% |
| | Total | 32% | | | | 44% | | 45% |

B

| OGp | Arabidopsis gene names | *At* genes | | | | *Bd* genes | | | | | *Hv* genes | | | |
|---|---|---|---|---|---|---|---|---|---|---|---|---|---|---|
| OGp0000529 | *LYSINE HISTIDINE TRANSPORTER 2 (LHT2); AMINO ACID TRANSPORTER-LIKE PROTEIN 2 (AATL2); (LHT1)* | | | -1.2 | | | | -0.94 | | | | | | -1.05 |
| OGp0000758 | *AMINO ACID PERMEASE 3 (AAP3) (AAP2) (AAP4)* | | -1.09 | -1.15 | | | -1.63 | | | | | | | -1.45 |
| OGp0003040 | *AMINO ACID TRANSPORTER 1 (AAT1)* | -2.18 | | | | -1.3 | | | | | | | -0.96 | |
| OGp0004800 | *Transmembrane amino acid transporter family protein (AT5G41800)* | -1.05 | | | | -1.01 | -2.42 | | | | -0.63 | -2.28 | | |
| OGp0012722 | *AMINO ACID PERMEASE 6 (AAP6)* | -1.24 | | | | -1.75 | | | | | 0.56 | | | |
| OGp0010057 | *Transmembrane amino acid transporter family protein (AT2G41190)* | 0.62 | | | | 0.9 | | | | | | | | |
| OGp0012770 | *GLUTAMINE DUMPER 3 (GDU3)* | -0.87 | | | | 1.31 | | | | | | | | |
| OGp0001664 | *CATIONIC AMINO ACID TRANSPORTER 8 (CAT8) (CAT5)* | -0.41 | | | | | | | | | -0.73 | -1.97 | | |
| OGp0000764 | *LIKE AUXIN RESISTANT 2 (LAX3) (LAX2); AUXIN RESISTANT 1 (AUX1); (LAX1)* | | -0.56 | | -0.54 | | | | | | | 0.37 | | |
| OGp0002730 | *Transmembrane amino acid transporter family protein (AVT3B) (AT4G38250) (AVT3)* | | | | | 0.79 | | | | | 1.18 | | | |
| OGp0009451 | *Transmembrane amino acid transporter family protein (AT1G25530)* | | | | | -0.9 | | | | | -0.8 | | | |
| **OGp0002892** | ***AROMATIC AND NEUTRAL TRANSPORTER 1 (ANT1)*** | | | | | 2.03 | | | | | | | -1.22 | |
| **OGp0002690** | ***Transmembrane amino acid transporter family protein (AT2G39130) (AT3G54830)*** | 0.81 | 8.5 | | | | | | | | | | | |
| OGp0004060 | *PROLINE TRANSPORTER 3 (PROT3) (PROT1) (PROT2)* | | -0.97 | | | | | | | | | | | |
| OGp0009517 | *Transmembrane amino acid transporter family protein (AT1G80510)* | -0.53 | | | | | | | | | | | | |
| OGp0003590 | *USUALLY MULTIPLE ACIDS MOVE IN AND OUT TRANSPORTERS 19 (UMAMIT19) (UMAMIT18) (UMAMIT17)* | -3.57 | 1.32 | -0.83 | | | | | | | | | | |
| OGp0004006 | *Transmembrane amino acid transporter family protein (AT2G40420) (AT3G56200)* | | | | | 1.15 | | | | | | | | |
| OGp0009061 | *MITOCHONDRIAL BASIC AMINO ACID CARRIER 2 (MBAC2)* | | | | | 1.44 | | | | | | | | |
| OGp0011133 | *PLANT UNCOUPLING MITOCHONDRIAL PROTEIN 1 (PUMP1)* | | | | | 0.89 | | | | | | | | |
| OGp0002889 | *Transmembrane amino acid transporter family protein (AT3G30390) (AT5G38820)* | | | | | -0.53 | | | | | | | | |
| OGp0007694 | *A BOUT DE SOUFFLE (BOU)* | | | | | -0.67 | | | | | | | | |
| OGp0009812 | *Transmembrane amino acid transporter family protein (AT1G08230)* | | | | | -0.65 | | | | | | | | |
| OGp0012188 | *Mitochondrial import inner membrane translocase subunit Tim17/Tim22/Tim23 family protein (ATOEP16-S)* | | | | | -1.22 | | | | | | | | |
| OGp0006617 | *BIDIRECTIONAL AMINO ACID TRANSPORTER 1 (BAT1)* | | | | | | | | | | | -0.85 | | |
| OGp0013238 | *AMINO ACID PERMEASE 7 (AAP7)* | | | | | | | | | | -0.35 | | | |
| OGp0006795 | *CATIONIC AMINO ACID TRANSPORTER 7 (CAT7)(CAT6)* | | | | | | | | | | | | | |
| OGp0009410 | *Transmembrane amino acid transporter family protein (AT1G47670)* | | | | | | | | | | | | | |
| OGp0006505 | *OUTER PLASTID ENVELOPE PROTEIN 16-L (ATOEP16-L)* | | | | | | | | | | | | | |
| OGp0010256 | *Tryptophan/tyrosine permease (AT2G33260)* | | | | | | | | | | | | | |
| OGp0010448 | *Mitochondrial substrate carrier family protein (MBAC1)* | | | | | | | | | | | | | |
| OGp0012453 | *Tryptophan/tyrosine permease (AT5G19500)* | | | | | | | | | | | | | |
| OGp0013170 | *DICARBOXYLATE TRANSPORT 2.1 (DIT2.1)* | | | | | | | | | | | | | |
| | Up | 8% | | | | 15% | | | | | 6% | | | |
| | Down | 28% | | | | 24% | | | | | 21% | | | |
| | Total | 36% | | | | 39% | | | | | 26% | | | |

**Supplementary Table 9: Gene nitrate-response in tri-species OGps belonging to selected significantly enriched nitrogen-related GO terms.** GO terms "cellular response to nitrogen starvation" [GO:0006995] **(A)** and "amino acid transmembrane transport" [GO:0003333] **(B)** were identified by the comparative analysis as specific of Arabidopsis-barley and Arabidopsis-Brachypodium subgroups, respectively. Genes associated to those GO terms and belonging to tri-species OGps were extracted. Lines represent OGps; colored rectangles represent genes, grouped by species (*At*, Arabidopsis; *Bd*, Brachypodium; *Hv*, barley). Color scale and numbers indicate maximum $Log_2FC$ values (1.5h or 3h nitrate *vs* mock); blue, upregulated; red, downregulated; grey, not-regulated/not-detected. OGps in bold are discussed in the text. Proportions of upregulated, downregulated and total DEGs among represented genes are indicated below the tables for each species.

| OGp | Arabidopsis gene identifiers | *At* genes | *Bd* genes | *Hv* genes |
|---|---|---|---|---|
| OGp0001036 | *AT2G38090, AT3G11280, AT5G01200, AT5G05790, AT5G58900* | -1.41 -1.17 1.47 | -2.30 0.57 | -0.94 |
| OGp0001053 | *AT2G22540, AT4G24540* | -0.53 | 0.55 | 0.56 1.01 |
| OGp0001638 | *AT1G14920, AT1G66350, AT2G01570, AT3G03450, AT5G17490* | -0.81 -0.87 | -0.93 | 0.57 |
| OGp0001976 | *AT3G46130, AT5G59780* | 1.03 0.95 | 0.82 | -0.88 -2.22 0.65 |
| OGp0007247 | *AT4G25420, AT5G51810* | -4.01 | -1.97 | 0.96 |
| OGp0000611 | *AT1G09540, AT1G57560* | -1.39 -1.53 | -0.72 | |
| OGp0006507 | *AT2G04240* | -0.83 | -0.80 -1.13 | |
| OGp0001236 | *AT1G49580, AT2G41140, AT3G19100, AT3G50530, AT3G56760* | 0.42 0.46 -0.42 | -0.43 | |
| OGp0001283 | *AT1G45050, AT1G75440, AT4G36410, AT5G42990* | 0.61 -0.71 1.38 | 0.51 | |
| OGp0011890 | *AT4G23060* | -1.12 | 0.55 | |
| OGp0013115 | *AT5G37260* | 2.94 | | 0.40 |
| OGp0000289 | *AT1G78290, AT3G50500, AT4G33950, AT4G40010, AT5G66880* | -0.68 | | -0.87 |
| OGp0000619 | *AT1G07540, AT3G12560, AT3G46590, AT5G13820, AT5G59430* | -0.61 -0.51 -0.76 | | -0.81 |
| OGp0003890 | *AT1G35515, AT4G09460* | -0.94 | | -0.92 |
| OGp0004682 | *AT4G20260, AT5G44610* | -0.84 | | -0.35 |
| OGp0005957 | *AT1G01060, AT2G46830* | -0.96 | | -0.42 |
| OGp0000772 | *AT1G26945, AT1G74500, AT3G28857, AT3G47710, AT5G15160, AT5G39860* | -1.12 | | 1.28 |
| OGp0006684 | *AT3G49690, AT5G65790* | -0.68 | | 0.61 |
| OGp0010455 | *AT2G43060* | -0.85 | | 0.40 |
| OGp0000543 | *AT4G25000* | | 1.10 | 1.69 |
| OGp0001968 | *AT3G10595, AT5G04760* | | 1.21 | 1.05 |
| OGp0000991 | *AT1G79460* | | -1.18 | -1.50 -0.83 -0.75 |
| OGp0004830 | *AT5G11260* | | -1.38 | -1.17 |
| OGp0007531 | *AT5G47390* | | 1.05 | -0.60 |
| OGp0001652 | *AT1G07430, AT2G29380, AT3G11410, AT5G59220* | 1.01 | | |
| OGp0002451 | *AT1G74670, AT3G02885, AT3G10185, AT5G15230* | 0.58 | | |
| OGp0010386 | *AT2G38560* | 0.44 | | |
| OGp0002618 | *AT1G67030, AT1G68360, AT5G06650* | -0.96 | | |
| OGp0002666 | *AT2G37630* | -0.64 | | |
| OGp0004299 | *AT3G05120, AT3G63010, AT5G27320* | -0.40 -1.35 | | |
| OGp0006141 | *AT1G69530, AT2G03090* | -0.78 | | |
| OGp0000948 | *AT1G18835, AT1G74660, AT1G75240, AT3G28917, AT5G42780* | -0.98 0.59 | | |
| OGp0001264 | *AT1G74650, AT3G28910, AT3G47600, AT5G62470* | -1.00 1.59 | | |
| OGp0001860 | *AT2G28950, AT2G37640, AT2G39700, AT3G55500, AT5G02260* | -0.48 1.05 | | |
| OGp0002720 | *AT2G45660, AT5G51860, AT5G51870, AT5G62165* | -0.65 0.61 | | |
| OGp0000647 | *AT1G06180, AT2G31180, AT3G23250* | | 0.72 | |
| OGp0003849 | *AT1G49010, AT5G08520, AT5G23650* | | 1.94 | |
| OGp0010536 | *AT2G36890* | | 0.57 | |
| OGp0012534 | *AT5G48170* | | 0.72 | |
| OGp0002038 | *AT4G02780* | | -5.12 | |
| OGp0003150 | *AT5G25900* | | -1.48 -1.64 -2.25 | |
| OGp0003825 | *AT1G01520, AT4G01280* | | -0.67 | |
| OGp0001826 | *AT1G18080, AT1G48630, AT3G18130* | | | 0.47 |
| OGp0003931 | *AT1G05710, AT2G31730* | | | 1.11 |
| OGp0006421 | *AT2G40830, AT3G56580* | | | 0.35 |
| OGp0010755 | *AT3G16350* | | | 0.48 |
| OGp0000665 | *AT2G23290, AT3G50060, AT4G37260, AT5G67300* | | | |
| OGp0000852 | *AT3G03940, AT5G18190* | | | |
| OGp0000776 | *AT1G19790, AT1G75520, AT2G18120, AT2G21400, AT3G51060, AT4G36260, AT5G33210, AT5G66350* | | | |
| OGp0001247 | *AT1G22690, AT1G75750, AT2G18420, AT4G09600, AT4G09610* | | | |
| OGp0001349 | *AT1G10588, AT2G39540, AT5G59845* | | | |
| OGp0001759 | *AT1G69600, AT3G28920, AT5G15210, AT5G39760, AT5G60480* | | | |
| OGp0002397 | *AT1G29640, AT2G34340, AT4G18980, AT5G45630* | | | |
| OGp0003621 | *AT1G26960, AT1G69780* | | | |
| OGp0004101 | *AT2G30810* | | | |
| OGp0004058 | *AT2G39880, AT3G09230, AT3G55730* | | | |
| OGp0004098 | *AT2G40740, AT3G56390* | | | |
| OGp0003921 | *AT1G28160, AT5G13910* | | | |
| OGp0004466 | *AT3G01140, AT5G15310* | | | |
| OGp0004818 | *AT5G50915* | | | |
| OGp0007282 | *AT4G30410, AT5G57780* | | | |
| OGp0009620 | *AT1G48270* | | | |
| OGp0009184 | *AT1G49950* | | | |
| OGp0009645 | *AT1G50420* | | | |
| OGp0010231 | *AT2G03500* | | | |
| OGp0009994 | *AT2G26300* | | | |
| OGp0010913 | *AT3G11540* | | | |
| OGp0012100 | *AT4G24210* | | | |
| OGp0012524 | *AT5G04010* | | | |
| OGp0012296 | *AT5G61850* | | | |
| | Up | 9% | 11% | 11% |
| | Down | 18% | 12% | 10% |
| | Total | 27% | 23% | 21% |

**Supplementary Table 10: Gene nitrate-response in tri-species OGps belonging to "response to gibberellin" GO term [GO:0009739] .** The GO term was identified as common to the 3 species by the comparative analysis on tri-species OGps. Lines represent OGps; colored rectangles represent genes, grouped by species (*At*, Arabidopsis; *Bd*, Brachypodium; *Hv*, barley). Color scale and numbers indicate maximum $Log_2FC$ values (1.5h or 3h nitrate *vs* mock); blue, upregulated; red, downregulated; grey, not-regulated/not-detected. OGps in bold are discussed in the text. Proportions of upregulated, downregulated and total DEGs among represented genes are indicated below the table for each species.

**Supplementary Table 11: Gene nitrate-response in tri-species OGps belonging to "rRNA processing" GO term [GO:0006364].** The GO term was identified as Arabidopsis-specific by the comparative analysis on tri-species OGps. Genes associated to this GO term and belonging to tri-species OGps were extracted. Lines represent OGps; colored rectangles represent genes, grouped by species (*At*, Arabidopsis; *Bd*, Brachypodium; *Hv*, barley). Color scale and numbers indicate maximum $Log_2FC$ values (1.5h or 3h nitrate *vs* mock); blue, upregulated; red, downregulated; grey, not-regulated/not-detected. Proportions for each species of upregulated, downregulated and total DEGs are indicated below the table.

| OGp | Arabidopsis gene names | *At* genes | *Bd* genes | *Hv* genes |
|---|---|---|---|---|
| OGp0007638 | *ROOT INITIATION DEFECTIVE 2 (RID2)* | 0.68 | 1.68 | |
| OGp0001082 | *Pre-rRNA-processing protein TSR2 (AT3G22510) (AT5G27990)* | 0.60 | | 0.36 |
| OGp0001099 | *Transducin/WD40 repeat-like superfamily protein (AT5G14050)* | 0.96 | | 0.54 |
| OGp0003185 | *Rrp15p protein (AT5G48240)* | 0.98 | | 0.41 |
| OGp0003886 | *Ribosomal protein L1p/L10e family (AT1G06380) (AT2G42650) (AT3G58660)* | 0.68 0.84 0.81 | | 0.43 |
| OGp0003821 | *Sas10/Utp3/C1D family (AT1G07840)* | 0.71 | | 0.53 |
| OGp0004384 | *HOMOLOGUE OF NAP57 (NAP57)* | 0.63 | | 0.59 0.49 |
| OGp0004616 | *FIBRILLARIN 2 (FIB2) (FIB1)* | 0.61 0.49 | | 0.45 |
| OGp0004836 | *RNA HELICASE10 (RH10)* | 0.57 | | 0.43 |
| OGp0006363 | *ARABIDOPSIS HOMOLOG OF YEAST RRS1 (ARRS1)* | 0.74 | | 0.34 |
| OGp0006543 | *EMBRYO DEFECTIVE (EMB2777) (EMB2796)* | 0.64 | | 0.56 |
| OGp0007180 | *YAOZHE (YAO); EMBRYO DEFECTIVE 2271 (EMB2271)* | 0.69 | | 0.45 |
| OGp0008724 | *RNA HELICASE 36 (RH36)* | 0.70 | | 0.41 |
| OGp0013030 | *GLUCOSE HYPERSENSITIVE 40 (GHS40)* | 0.73 | | 0.97 |
| OGp0001252 | *INVOLVED IN RRNA PROCESSING 5 (IRP5) (IRP7)* | 0.45 | | |
| OGp0001744 | *Ribosomal protein L1p/L10e family (AT1G08360) ; PIGGYBACK1 (PGY1); (AT5G22440)* | 0.41 | | |
| OGp0001798 | *NUCLEOLIN LIKE 1 (NUC-L1) (NUC-L2) (AT4G08760)* | 0.72 1.20 | | |
| OGp0001997 | *Transducin/WD40 repeat-like superfamily protein (AT3G10530) (AT3G23850)* | 0.87 | | |
| OGp0002652 | *RRP6-LIKE (RRP6L1) (RRP6L2)* | 0.40 | | |
| OGp0002745 | *BLOCK OF CELL PROLIFERATION 1 (BOP1)* | 0.61 | | |
| OGp0002935 | *P-loop containing nucleoside triphosphate hydrolases superfamily protein (AT3G18600)* | 0.85 | | |
| OGp0003043 | *LA PROTEIN 1 (LA1)* | 0.55 | | |
| OGp0003791 | *ARABIDOPSIS HOMOLOGUE OF YEAST BRX1 (ATBRX1-2) (ATBRX1-1)* | 0.61 0.74 | | |
| OGp0004081 | *CHLOROPLAST RNA-BINDING PROTEIN 29 (AT2G37220) (CP29)* | 0.81 | | |
| OGp0004607 | *MATERNAL EFFECT EMBRYO ARREST 49 (MEE49)* | 0.67 | | |
| OGp0004658 | *U3 ribonucleoprotein (Utp) family protein (AT4G02400) (AT5G08600) (AT5G36980)* | 0.81 | | |
| OGp0005548 | *RIBOSOMAL RNA PROCESSING 4 (RRP4)* | 0.60 | | |
| OGp0006091 | *rRNA-processing EFG1-like protein (DUF2361) (AT1G04230) (AT5G43720)* | 0.56 | | |
| OGp0005769 | *YEAST LSG1 ORTHOLOGUE (LSG1-2) (LSG1-1)* | 0.75 | | |
| OGp0005937 | *GNAT acetyltransferase (DUF699) (AT1G10490) (AT3G57940)* | 0.47 0.84 | | |
| OGp0006033 | *LANT-SPECIFIC COMPONENT OF THE PRE- RRNA PROCESSING COMPLEX1 (PCP1)* | 0.66 | | |
| OGp0005767 | *LANT-SPECIFIC COMPONENT OF THE PRE- RRNA PROCESSING COMPLEX2 (PCP2)* | 0.59 | | |
| OGp0005853 | *TRNA METHYLTRANSFERASE 112B (TRM112B) (ATTRM112A)* | 0.49 | | |
| OGp0005935 | *ESSENTIAL NUCLEAR PROTEIN 1 (ENP1)* | 0.78 | | |
| OGp0005758 | *Chloroplast Mini-RNase III-like enzymes (RNC3) (RNC4)* | 1.03 | | |
| OGp0005692 | *Nucleolar protein (AT1G63810)* | 0.75 | | |
| OGp0005876 | *HARBINGER TRANSPOSON-DERIVED PROTEIN 1 (HDP1); IDM1-ASSOCIATED PROTEIN 1 (IDAP1)* | 0.51 | | |
| OGp0004161 | *Ribosome biogenesis protein (AT2G01640)* | 0.47 | | |
| OGp0006440 | *PNAS-3-like protein (AT2G25355) (AT4G32175)* | 0.51 | | |
| OGp0006549 | *PIGMENT DEFECTIVE 328 (PDE328); Pseudouridine synthase family protein (AT3G43340)* | 0.67 | | |
| OGp0007149 | *ARM repeat superfamily protein (AT3G06530)* | 0.81 | | |
| OGp0006856 | *RIBOSOMAL RNA PROCESSING 5 (RRP5)* | 0.68 | | |
| OGp0006999 | *ARABIDOPSIS HOMOLOG OF YEAST RPF2 (ARPF2)* | 0.82 | | |
| OGp0006873 | *KRR1 family protein (AT3G24080)* | 0.75 | | |
| OGp0006582 | *Exosome complex exonuclease RRP46-like protein ; Ribosomal protein S5 domain 2-like superfamily protein* | 0.61 | | |
| OGp0007016 | *ROOT INITIATION DEFECTIVE 3 (RID3)* | 0.56 | | |
| OGp0006708 | *RNA-binding (RRM/RBD/RNP motifs) family protein (AT3G56510)* | 0.45 | | |
| OGp0006966 | *Nucleolar essential protein-like protein (AT3G57000)* | 0.50 | | |
| OGp0007020 | *U3 SMALL NUCLEOLAR RNA-ASSOCIATED PROTEIN 11 (UTP11)* | 0.60 | | |
| OGp0007230 | *TRNA METHYLTRANSFERASE 4D (TRM4D) (TRM4C)* | 0.97 0.84 | | |
| OGp0007753 | *RNA-binding KH domain-containing protein (AT5G08420)* | 0.53 | | |
| OGp0007735 | *PESCADILLO (PES)* | 0.59 | | |
| OGp0007579 | *RNA cyclase family protein (AT5G22100)* | 0.81 | | |
| OGp0007470 | *U3 small nucleolar ribonucleoprotein (AT5G66540)* | 0.87 | | |
| OGp0009609 | *P-loop containing nucleoside triphosphate hydrolases superfamily protein (AT1G06720)* | 0.76 | | |
| OGp0008801 | *CRUCIFERIN B (CRB)* | 0.95 | | |
| OGp0008669 | *rRNA biogenesis RRP36-like protein (AT1G12650)* | 0.69 | | |
| OGp0009567 | *PERIODIC TRYPTOPHAN PROTEIN 2 (PWP2)* | 0.65 | | |
| OGp0009697 | *NUCLEOLAR FACTOR 1 (NOF1)* | 0.57 | | |
| OGp0008866 | *MAK16 protein-like protein* | 0.87 | | |
| OGp0009120 | *REDUCED POLLEN NUMBER 1 (RDP1)* | 0.60 | | |
| OGp0008988 | *STRESS RESPONSE SUPPRESSOR 1 (STRS1)* | 0.57 | | |
| OGp0009514 | *FASCIATED STEM 4 (FAS4)* | 0.64 | | |
| OGp0008683 | *pre-rRNA-processing TSR1-like protein (AT1G42440)* | 0.61 | | |
| OGp0008846 | *HOMOLOG OF YEAST MTR4 (MTR4)* | 0.38 | | |
| OGp0008945 | *3'-5'-exoribonuclease family protein (AT1G60080)* | 0.61 | | |
| OGp0008759 | *Ribosomal RNA processing Brix domain protein (IMP4)* | 0.66 | | |
| OGp0009057 | *DEAD-BOX RNA HELICASE 29 (RH29)* | 0.79 | | |
| OGp0010404 | *NUCLEOLAR COMPLEX ASSOCIATED 4 (NOC4)* | 0.75 | | |
| OGp0009925 | *Zinc ion binding protein (AT2G19385)* | 0.62 | | |
| OGp0010334 | *YBEY ENDORIBONUCLEASE (ATYBEY)* | 0.55 | | |
| OGp0010388 | *HUMAN WDR55 (WD40 REPEAT) HOMOLOG (WDR55)* | 0.73 | | |
| OGp0010587 | *Radical SAM superfamily protein (AT2G39670)* | 0.53 | | |
| OGp0010641 | *SMALL ORGAN 4 (SMO4)* | 0.77 | | |
| OGp0010085 | *Ribosomal protein L1p/L10e family (AT2G42710)* | 0.51 | | |
| OGp0010038 | *Axoneme-associated protein MST101(2) protein (AT2G44820)* | 0.58 | | |
| OGp0011270 | *pre-rRNA-processing ESF1-like protein (AT3G01160)* | 0.76 | | |
| OGp0011276 | *Polynucleotidyl transferase, ribonuclease H-like superfamily protein (AT3G15080)* | 0.81 | | |
| OGp0011304 | *Transducin family protein / WD-40 repeat family protein (AT3G21540)* | 0.73 | | |
| OGp0011253 | *rRNA processing protein-like protein (EBP2)* | 0.92 | | |
| OGp0011284 | *Protein serine/threonine kinase (AT3G51270)* | 0.48 | | |
| OGp0011498 | *ERBB-3 BINDING PROTEIN 1 (EBP1)* | 0.55 | | |
| OGp0010718 | *PIGMENT DEFECTIVE 322 (PDE322)* | 0.62 | | |
| OGp0011213 | *EUKARYOTIC INITIATION FACOR 6A (eIF6A)* | 0.44 | | |
| OGp0011180 | *EMBRYO SAC DEVELOPMENT ARREST 7 (EDA7)* | 0.56 | | |
| OGp0011244 | *3'-5'-exoribonuclease family protein (RRP41)* | 0.43 | | |
| OGp0011995 | *transducin family protein / WD-40 repeat family protein (AT4G04940)* | 0.70 | | |
| OGp0011594 | *Ribosomal protein L7Ae/L30e/S12e/Gadd45 family protein (AT4G22380)* | 0.50 | | |
| OGp0011945 | *TRNA METHYLTRANSFERASE 7B (TRM7B)* | 0.86 | | |
| OGp0011694 | *U3 small nucleolar RNA-associated-like protein (AT4G28200)* | 0.45 | | |
| OGp0011789 | *WD repeat and SOF domain-containing protein 1 (AT4G28450)* | 0.73 | | |
| OGp0012374 | *Ribosomal protein S8e family protein (AT5G06360)* | 0.49 | | |
| OGp0012392 | *Ribosomal protein L7Ae/L30e/S12e/Gadd45 family protein (AT5G08180)* | 0.44 | | |
| OGp0013308 | *polyribonucleotide nucleotidyltransferase (AT5G14580)* | 0.55 | | |
| OGp0013204 | *PESCADILLO ORTHOLOG (PEP2)* | 0.55 | | |
| OGp0013311 | *Alpha-L RNA-binding motif/Ribosomal protein S4 family protein (AT5G15750)* | 0.69 | | |
| OGp0012901 | *TORMOZEMBRYO DEFECTIVE (TOZ)* | 0.67 | | |
| OGp0013193 | *Ribosomal RNA processing-like protein (AT5G20600)* | 0.51 | | |
| OGp0012977 | *RIBOSOMAL RNA PROCESSING7 I (RRP7)* | 0.88 | | |
| OGp0012580 | *S-adenosyl-L-methionine-dependent methyltransferases superfamily protein (AT5G40530)* | 0.74 | | |
| OGp0012648 | *RNA-binding NOB1-like protein (NOB1)* | 0.54 | | |
| OGp0004066 | *Ribosomal protein L7Ae/L30e/S12e/Gadd45 family protein (AT2G47610) (AT3G62870)* | | 0.46 | |
| OGp0005816 | *DCL protein (DUF3223) (AT1G45230)* | | 3.01 | |
| OGp0010524 | *PLASTID RIBOSOMAL PROTEIN OF THE 30S SUBUNIT 5 (PRPS5)* | | 0.76 | |
| OGp0001398 | *Ribosomal protein S5 domain 2-like superfamily protein (AT2G09990) (AT3G04230) (AT5G18380)* | | | 0.53 0.40 |
| OGp0006827 | *DCL protein (DUF3223) (AT3G46630)* | | | 1.14 |
| OGp0006695 | *EVERSHED1 (EVR1)* | | | 0.35 |
| OGp0009970 | *SLOW WALKER1 (SWA1)* | | | 0.46 |
| OGp0000229 | *RIBOSOMAL PROTEIN S11 (RPS11)* | | | |
| OGp0000833 | *GROWTH-REGULATING FACTOR 3 (GRF3) (GRF4)* | | | |
| OGp0001933 | *Ribosomal protein S11 family protein* | | | |
| OGp0001971 | *RIBOSOMAL PROTEIN S28 (RPS28)* | | | |
| OGp0002810 | *Ribosomal protein L30/L7 family protein (RPL7B)* | | | |
| OGp0002722 | *ADENOSINE DIMETHYL TRANSFERASE 1A (DIM1A)* | | | |
| OGp0002906 | *RNA HELICASE 14 (RH14) (RH46)* | | | |
| OGp0002886 | *RNASE THREE-LIKE PROTEIN 2 (RTL2) (RTL3)* | | | |
| OGp0003144 | *MERISTEM-DEFECTIVE (MDF)* | | | |
| OGp0003895 | *MULTIPLE ORGANELLAR RNA EDITING FACTOR 5 (MORF5) (MORF6)* | | | |
| OGp0003790 | *40S RIBOSOMAL PROTEIN SA (P40)* | | | |
| OGp0004135 | *Ribosomal L29 family protein* | | | |
| OGp0004103 | *SIMILAR TO YEAST POP4 (POP4)* | | | |
| OGp0003996 | *SM-LIKE 6 (LSM6B) (LSM6A)* | | | |
| OGp0004332 | *INVOLVED IN RRNA PROCESSING 1 (IRP1)* | | | |
| OGp0004513 | *Polynucleotidyl transferase, ribonuclease H-like superfamily protein* | | | |
| OGp0004823 | *Ribosomal protein S8e family protein* | | | |
| OGp0005730 | *EMBRYO DEFECTIVE 1687 (EMB1687)* | | | |
| OGp0006088 | *RHO-N 1 (RHON1)* | | | |
| OGp0005612 | *INVOLVED IN RRNA PROCESSING 8 (IRP8); INTERACTING WITH DNA-BINDING DOMAIN OF ZN-FINGER PARP 1 (DIP2)* | | | |
| OGp0005903 | *5'-3' EXORIBONUCLEASE 3 (XRN3)* | | | |
| OGp0006282 | *HUA ENHANCER 2 (HEN2)* | | | |
| OGp0006295 | *(RAP)* | | | |
| OGp0006953 | *Ribosomal protein S7e family protein* | | | |
| OGp0007074 | *H/ACA ribonucleoprotein complex, subunit Gar1/Naf1 protein* | | | |
| OGp0006766 | *RIBOSOMAL RNA PROCESSING 42 (RRP42)* | | | |
| OGp0006959 | *INVOLVED IN RRNA PROCESSING 4 (IRP4); ASYMMETRIC LEAVES ENHANCER 3 (AE3)* | | | |
| OGp0006875 | *RIBONUCLEASE PH45A (RRP45a); ECERIFERUM 7 (CER7)* | | | |
| OGp0006914 | *Exonuclease family protein* | | | |
| OGp0007163 | *MORPHOLOGY OF AGO1-52 SUPPRESSED2 (MAS2)* | | | |
| OGp0007347 | *INVOLVED IN RRNA PROCESSING 3 (IRP3/BPL1) ; BPA1-LIKE 5 (BPL5)* | | | |
| OGp0007188 | *31-KDA RNA BINDING PROTEIN (RBP31); CHLOROPLAST RNA-BINDING PROTEIN 31B (CP31B)* | | | |
| OGp0007540 | *Methyltransferase* | | | |
| OGp0007771 | *Scarecrow-like transcription factor 11 (SCL11)* | | | |
| OGp0009818 | *RNA-binding (RRM/RBD/RNP motifs) family protein* | | | |
| OGp0009165 | *PALEFACE 1 (PFC1)* | | | |
| OGp0009028 | *Ribosomal RNA-processing protein* | | | |
| OGp0009813 | *Polynucleotidyl transferase, ribonuclease H-like superfamily protein* | | | |
| OGp0008780 | *S1 DOMAIN-CONTAINING RBP (SDP)* | | | |
| OGp0009228 | *CLP PROTEASE PROTEOLYTIC SUBUNIT 1 (CLPR1)* | | | |
| OGp0009496 | *Methyltransferase* | | | |
| OGp0009389 | *TARGET OF RAPAMYCIN (TOR)* | | | |
| OGp0009168 | *RNA HELICASE 20 (RH20)* | | | |
| OGp0009722 | *Radical SAM superfamily protein* | | | |
| OGp0009588 | *MIDASIN 1 (MDN1)* | | | |
| OGp0009240 | *HIGH PHOTOSYNTHETIC EFFICIENCY 1 (HPE1)* | | | |
| OGp0008715 | *RIBOSOMAL PROTEIN S9 (RPS9)* | | | |
| OGp0009811 | *Pseudouridine synthase family protein* | | | |
| OGp0008744 | *RRP44 HOMOLOG B (RRP44B)* | | | |
| OGp0010679 | *RNASE E/G-LIKE (RNEE/G)* | | | |
| OGp0010659 | *EMBRYO DEFECTIVE 2763 (EMB2763)* | | | |
| OGp0010413 | *EMBRYO SAC DEVELOPMENT ARREST 27 (EDA27)* | | | |
| OGp0010672 | *FIONA1 (FIO1)* | | | |
| OGp0010492 | *RRP6-LIKE 3 (RRP6L3)* | | | |
| OGp0010914 | *POLYNUCLEOTIDE PHOSPHORYLASE (PNP)* | | | |
| OGp0011492 | *TRNA METHYLTRANSFERASE 4E (TRM4E)* | | | |
| OGp0011365 | *ENHANCER OF RNAI (ERI-1)* | | | |
| OGp0011243 | *Radical SAM superfamily protein* | | | |
| OGp0011357 | *S1 RNA-BINDING RIBOSOMAL PROTEIN 1 (SRRP1)* | | | |
| OGp0011002 | *RIBOSOMAL PROTEIN S9 M (RPS9M)* | | | |
| OGp0011282 | *PLASTID-SPECIFIC RIBOSOMAL PROTEIN 2 (PSRP2)* | | | |
| OGp0011271 | *CHLOROPLAST STEM-LOOP BINDING PROTEIN OF 41 KDA (CSP41A)* | | | |
| OGp0011196 | *EMBRYO DEFECTIVE 3126 (EMB3126)* | | | |
| OGp0012091 | *P-loop containing nucleoside triphosphate hydrolases superfamily protein* | | | |
| OGp0011877 | *HIT-type Zinc finger family protein* | | | |
| OGp0012054 | *RBFA DOMAIN-CONTAINING PROTEIN 1 (RBF1)* | | | |
| OGp0012531 | *EMBRYO DEFECTIVE 2730 (EMB2730)* | | | |
| OGp0013060 | *CHLOROPLAST MRAW-LIKE (CMAL)* | | | |
| OGp0013359 | *Pre-mRNA cleavage complex II protein family (GRC3)* | | | |
| OGp0013350 | *SUPPRESSOR OF VARIEGATION 3 (SVR3)* | | | |
| OGp0012883 | *TRNA METHYLTRANSFERASE 7C (TRM7C)* | | | |
| OGp0012521 | *EMBRYO DEFECTIVE 269 (EMB269)* | | | |
| OGp0012544 | *TRNA METHYLTRANSFERASE 4H (TRM4H)* | | | |
| OGp0012674 | *16S rRNA processing protein RimM family* | | | |
| OGp0013444 | *Pseudouridine synthase family protein* | | | |
| OGp0013302 | *TRF4/5-LIKE (TRL)* | | | |
| OGp0013405 | *GAMETOPHYTE DEFECTIVE 1 (GAF1)* | | | |
| OGp0012908 | *EMBRYO DEFECTIVE 2746 (EMB2746)* | | | |
| OGp0013335 | *ADENOSINE DIMETHYL TRANSFERASE 1B (DIM1B)* | | | |
| OGp0012906 | *GTP-binding protein Era-like protein (ERA-1)* | | | |
| | Up | 45% | 2% | 7% |
| | Down | 0% | 0% | 0% |
| | Total | 45% | 2% | 7% |

A

| OGp | Arabidopsis gene names | *At* genes | *Bd* genes | *Hv* genes |
|---|---|---|---|---|
| OGp0008929 | *RAB GTPASE HOMOLOG G3B (RABG3B)* | -0.88 | 0.68 | |
| OGp0005976 | *RAB GTPASE HOMOLOG A3 (RABA3)* | -0.72 | | -0.81 |
| OGp0005445 | *RAB GTPASE HOMOLOG A2 (RABA2B) (RABA2C)* | | 0.69 | 0.55 |
| OGp0002639 | *RAB GTPASE HOMOLOG 5 (RABA5E) (RABA5D) (RABA5C) (RABA5B)* | -0.62 | | |
| OGp0004769 | *Ras-related small GTP-binding family protein (SGP1)* | -0.81 | | |
| OGp0006818 | *RAB GTPASE HOMOLOG C (RABC2B) (RABC2A)* | -1.09, -0.51 | | |
| OGp0000249 | *ARABIDOPSIS THALIANA RAB GTPASE HOMOLOG A1A (RABA1A) (RABA1B) (RABA1I) (RABA1H) (RABA1G) (RABA1E) (RABA1D) (RABA1C) (RABA1F)* | 0.71, 0.35, -0.94 | | |
| OGp0001966 | *RAB GTPASE HOMOLOG A4 (RABA4D) (RABA4B) (RABA4C) (RABA4A)* | | 0.76 | |
| OGp0004539 | *RAB GTPASE HOMOLOG B1B (RAB-B1B)* | | 0.49 | |
| OGp0005410 | *RAB GTPASE HOMOLOG B18 (RAB18)* | | 0.44 | |
| OGp0006756 | *RAB GTPASE HOMOLOG (RAB5C)* | | 0.45 | |
| OGp0013090 | *RAB GTPASE HOMOLOG A2D (RABA2D)* | | 0.93 | |
| OGp0000976 | *RAB GTPASE HOMOLOG G (RABG3E) (RABG3D) (RABG3C) (RABG1F)* | | | |
| OGp0001062 | *RAB GTPASE HOMOLOG 8 (RAB8A) (RAB8) (RAB8B) (RAB8C)* | | | |
| OGp0001734 | *RAB GTPASE HOMOLOG (RAB1C) (RABD2C) (RABD2B)* | | | |
| OGp0004042 | *RAB GTPASE HOMOLOG (RABH1D) (RABH1B) (RABH1E)* | | | |
| OGp0004602 | *RAB GTPASE HOMOLOG (ATRAB5B) (RAB5A)* | | | |
| OGp0008677 | *RAB GTPASE 11C (ATRAB11C)* | | | |
| OGp0012485 | *RAB GTPASE HOMOLOG A5A (RABA5A)* | | | |
| OGp0007687 | *LIGHT INSENSITIVE PERIOD1 (LIP1)* | | | |
| | Up | 4% | 21% | 3% |
| | Down | 15% | 0% | 3% |
| | Total | 19% | 21% | 7% |

B

| OGp | Arabidopsis gene names | *At* genes | *Bd* genes | *Hv* genes |
|---|---|---|---|---|
| OGp0009715 | *ECTOPIC ROOT HAIR 1 (ERH1)* | -0.51 | -0.63 | |
| OGp0001388 | *HOMOLOG OF CARROT EP3-3 CHITINASE (AT2G43580) (AT2G43590) (AT3G47540) (EP3)* | -0.98 | | |
| OGp0001442 | *PATHOGENESIS-RELATED 3 (PR3)* | -0.90 | | |
| OGp0012918 | *ENDO-BETA-MANNASE 7 (MAN7)* | -0.97 | | |
| OGp0010938 | *XYLOGLUCAN ENDO-TRANSGLYCOSYLASE-RELATED 8 (XTR8/XTH31)* | -0.42 | | |
| OGp0003839 | *Chitinase family protein (AT1G02360) (AT4G01700)* | | -1.57 | |
| OGp0010553 | *XYLOGLUCAN ENDOTRANSGLUCOSYLASE/HYDROLASE 32 (XTH32)* | | | -0.57 |
| OGp0006223 | *ENDO-BETA-MANNASE 2 (MAN2) (MAN5)* | | | |
| OGp0007482 | *ENDO-BETA-MANNASE 6 (MAN6)* | | | |
| | Up | 7% | 0% | 0% |
| | Down | 36% | 14% | 8% |
| | Total | 36% | 14% | 8% |

C

| OGp | Arabidopsis gene names | *At* genes | *Bd* genes | *Hv* genes |
|---|---|---|---|---|
| OGp0002576 | *UDP-GLUCOSE 4-EPIMERASE (UGE4/REB1) (AT3G28530) (UGE5) (UGE2)* | -0.69, -0.63 | 0.60 | -1.17 |
| OGp0001768 | *XYLOGLUCAN XYLOSYLTRANSFERASE (XXT4) (XXT5) (XXT3)* | -0.47 | | |
| OGp0006824 | *XYG XYLOSYLTRANSFERASE 1 (XXT1) (XXT2)* | -0.70, -0.51 | | |
| OGp0007614 | *XYLOGLUCAN L-SIDE CHAIN GALACTOSYLTRANSFERASE POSITION 2 (XLT2/GT18)* | -1.21 | | |
| OGp0010488 | *SHORT ROOT IN SALT MEDIUM 3 (RSA3/MUR3)* | | | 1.19 |
| OGp0001493 | *Glycosyl hydrolase family protein (BGLC1)* | | | |
| OGp0010030 | *FUCOSYLTRANSFERASE 1 (FUT1/MUR2)* | | | |
| | Up | 0% | 10% | 8% |
| | Down | 46% | 0% | 8% |
| | Total | 46% | 10% | 17% |

**Supplementary Table 12: Gene nitrate-response in tri-species OGps belonging to selected significantly enriched membrane trafficking and cell wall biogenesis GO terms.** GO terms “Rab protein signal transduction” [GO:0032482] **(A)**, “cell wall macromolecule catabolic process” [GO:0016998] **(B)** and “xyloglucan biosynthetic process” [GO:0009969] **(C)** were identified by the comparative analysis as specific of Brachypodium (upregulated genes), Arabidopsis-Brachypodium (downregulated genes) and Arabidopsis (downregulated genes), respectively. Genes associated to those GO terms and belonging to tri-species OGps were extracted. Lines represent OGps; colored rectangles represent genes, grouped by species (*At*, Arabidopsis; *Bd*, Brachypodium; *Hv*, barley). Color scale and numbers indicate maximum Log2FC values (1.5h or 3h nitrate vs mock); blue, upregulated; red, downregulated; purple, downregulated at 1.5h / upregulated at 3h; grey, not-regulated/not-detected. Proportions of upregulated, downregulated and total DEGs among represented genes are indicated below the tables for each species.
